# microRNA profile of endometrial cancer from Indian patients-Identification of potential biomarkers for prognosis

**DOI:** 10.1101/2023.08.07.552324

**Authors:** Shraddha Hegde, Kalpesh Wagh, Apoorva Abikar, Sughosha Nambiar, Shriraksha Ananthamurthy, Navyashree Hosahalli Narayana, Pallavi Venkateshaiah Reddihalli, Savitha Chandraiah, Suma Mysore Narayana, Prathibha Ranganathan

## Abstract

Endometrial cancer is one of the major cancers in women throughout the world. If diagnosed early, these cancers are treatable and the prognosis is usually good. However, one major problem in treating endometrial cancer is accurate diagnosis and staging. Till date, the choice method for diagnosis and staging is histopathology. Although there are few molecular markers identified, they are not always sufficient in making accurate diagnosis and deciding on therapeutic strategy. As a result, very often patients are under treated or over treated. In this study, our group has profiled microRNAs (miRNA) from Indian patients using NGS-based approach. We have identified differentially expressed microRNAs in endometrial cancer. These microRNAs have also been compared to data from TCGA (The Cancer Genome Atlas), which represent other populations and also correlated to relevance in overall survival. Using *in-silico* approaches, mRNA targets of the miRNAs have been predicted. After comparing with TCGA, we have identified 16 miRNA-mRNA pairs which could be potential prognostic biomarkers for endometrial cancer. This is the first miRNA profiling report from Indian cohort and one of the very few studies which have identified potential biomarkers of prognosis in endometrial cancer.

## Introduction

Endometrial cancer, arising from the lining of the uterus is the most common cancer of the female reproductive organs in the United States. It is estimated by The American Cancer Society that in 2023, there would be about 66,200 new cases and 13,030 women would die of this cancer. About 90% of the cancers of endometrium are carcinomas. 5 year survival rate for these cancers is 81% (1). Endometrial cancers are broadly classified as Type 1 lesions, which are the most common and are usually hormone sensitive and have good prognosis. These tumors are usually low-grade and have a background of hyperplasia. Type 2 lesions are higher grade and have higher recurrence frequencies (2). Most endometrial cancers present with symptoms such as vaginal bleeding and pelvic pain. These cancers are usually treatable if diagnosed early. For a long time, histopathology was the choice method for staging endometrial cancer and also for deciding on the course of therapy. However, studies have shown that cancers of the same stage and histology can have very distinct molecular and genomic profiles (reviewed in (3)). Identification of such molecular markers would aid in developing more effective and personalized therapies. The use of immune checkpoint inhibitor, pembrolizumab (anti-PD-1), in tumors with defective DNA mismatch repair would be beneficial to approximately 20-30% of patients with advanced endometrial cancer (4, 5) and,(reviewed in(3)). Other genomic changes and molecular markers in endometrial cancer, such as hormone receptor status, could lead to more tailored therapy in the future.

Several molecular prognostic biomarkers have been studied in terms of survival and therapeutic response in women with endometrial cancer. This includes hormone receptors, microRNAs (miRNA), and other molecules (Ex: HER2 (6), p21(7), HE4 (8), PTEN (9), p27 (10), ANCCA (11), and ANXA2 (12)), which have emerged as potentially useful biomarkers (reviewed in (13)). However, for a better understanding and classification of the disease and for making therapeutic decisions, it is essential to identify and critically evaluate more biomarkers and their relevance in prognosis.

miRNAs have been reported to have a substantial association with many diseases including cancer. miRNAs, particularly circulating and exosomal miRNAs have a huge potential to be used as diagnostic markers (reviewed in (14)). miRNAs have been shown to be stable in formalin-fixed, paraffin-embedded (FFPE) tissues in multiple cancer types. Also, miRNAs can be isolated and analyzed from body fluids such as serum, pancreatic juice, urine etc., which makes it possible to be analyzed from liquid biopsies. Differential miRNA profiles have been seen in pancreatic juice of chronic pancreatitis vs pancreatic adenocarcinoma patients (15). Differential profiles of miRNA have been seen in sera of patients with malignant and non-malignant lung disease (16). Analysis of urine samples have identified miRNAs which are potential diagnostic and prognostic biomarkers (17, 18). Besides cancer, miRNAs have been found to be differentially expressed in several other disease conditions. miRNAs are expressed in cell-type specific manner and due to the feasibility of analyzing these with non-invasive procedures, have a huge potential to be developed as biomarkers.

In cases of endometrial cancer, the classification is based mostly on histopathology. The major drawback of this approach is inter-observer variability and this substantially influences the therapeutic approaches leading to under or over treatment. Molecular markers would overcome this problem to some extent. Based on genomic abnormalities, TCGA has identified 4 major molecular subtypes namely POLE (DNA polymerase epsilon catalytic subunit) mutated, mismatch repair (MMR)–deficient, p53 abnormal, and no specific molecular profile (19). This requires a combination of immunohistochemical tests and mutation analysis. The efficacy of these molecular markers in a diagnostic/prognostic test needs evaluation (20).

There have been a few studies which have profiled miRNAs in endometrial cancer (21), compared different stages and sub-types (22) and also compared data available from TCGA to identify markers relevant in prognosis (23). However, all of the above studies are done on populations restricted to one region or data from TCGA. The study shown here presents data from Indian population which is mostly not represented even in public databases. The data has also been compared to TCGA data, which shows relevance in other populations.

## Materials and methods

### Collection of clinical samples

Samples used for this study have been collected from Kidwai Memorial Institute of Oncology (KMIO), Bengaluru and Vanivilas Hospital, Bengaluru, between the period of 2018-2022. This study was approved by the Institutional Ethics Committee at Kidwai Memorial Institute of Oncology, Vanivilas Hospital and Centre for Human Genetics where most of the molecular work was carried out. Women diagnosed with pathological stage I-IV endometroid type of endometrial carcinoma at KMIO between 2018 and 2022 were included in this study. The control samples were collected from patients undergoing hysterectomy for non-neoplastic conditions. Upon receiving the samples at the histopathology laboratory at KMIO, Hematoxylin and Eosin-stained sections was examined by the pathologist. Immunohistochemistry was performed wherever necessary to arrive at the diagnosis. Based on the histopathology report, only endometroid carcinoma samples along with controls were taken for miRNA profiling. Any other histology like polyps, hyperplasia and stromal tumors were rejected.

### RNA Isolation

All tissue samples were collected in RNA*later* (Sigma Life Sciences) and stored at −80^0^C till further processing. Before isolation of RNA, the tissues were rinsed with PBS and homogenized with RLT Buffer (QIAGEN GmbH) using a hand-held homogenizer. RNA was isolated using the Qiagen RNeasy Mini Kit (QIAGEN GmbH) according to manufacturer’s instructions. On column DNase digestion was also performed for all samples. The isolated RNA was quantified using NanoDrop (Thermo Scientific, USA).

### RNA – Sequencing and analysis

The RNA-sequencing was outsourced to Molsys Pvt Ltd, Bengaluru and Nucleome Informatics Pvt Ltd, Hyderabad.

44 small RNA libraries generated using NEB NextR Multiplex Small RNA Library Prep Set for IlluminaR were sequenced and the resultant fastq files showed the phred score greater than 30 for all samples. The reads were trimmed using the adapter sequence (AGATCGGAAGAGCACACGTCT). Reads that mapped to rRNA sequences were removed using Bowtie (24). Reads were size-selected for 18-25bp long reads using BBTools (25). Obtained sequences were then collapsed using the mapper tool from miRDeep2 (26) and used for identification and quantification of miRNA. DESeq2 function in R (27) was used to identify the differentially regulated miRNAs in endometrial cancer vs control samples. Differentially expressed novel miRNAs were identified at FDR<0.001 and four-fold change in expression between endometrial cancer samples versus control samples.

### Retrieval of data from The Cancer Genome Atlas (TCGA)

RNA-Seq data on endometrial cancer was retrieved from The Cancer Genome Atlas (28) database using the Xena Browser (29). The data from TCGA Endometrioid Cancer (UCEC) containing 26 datasets was selected for our study. This dataset had miRNA sequenced from 429 patient samples using IlluminaHiSeq miRNASeq. Information about 2238 sequenced miRNAs was obtained from TCGA for our analysis.

### Comparison of data obtained from miRNA seq and TCGA

Differential expression data obtained from our cohort (Indian population) identifies the miRNA using gene ID, whereas the data from TCGA identifies using accession numbers. In order to make the comparison, the first step was to have uniform identifiers. This was done as described in (30). The differentially expressed miRNAs were compared to TCGA data to obtain a common list of miRNAs which could be then used for further analysis such as survival analysis and target prediction.

### Survival Analysis

The TCGA Endometrioid Cancer (UCEC) cohort had a survival dataset for 583 patients in total. The survival data has endpoints such as Overall survival (OS), Progression-free interval (PFI), Disease-specific survival (DSS), Disease-free interval (DFI). The survival data was retrieved from TCGA and compared with the patients for which miRNA expression data was also available. 398 patients had both the relevant data and hence only these were used for further analysis.

For prognostic relevance of each miRNA, the patients were grouped into high expressers and low expressers based on the median expression value (as per TCGA data) and the survival probability was plotted against time using K-M plotter (31) (32) (33).

If the Hazard ratio was >1, higher expression of the miRNA corresponds to greater risk/lower survival; and if the Hazard ratio <1, higher expression corresponds to less risk/better survival.

### Statistical analysis

All the statistical analyses were performed using the R statistical package. In all the statistical tests, p < 0.05 was considered significant (34). Hazard ratio was also taken into consideration for the selection of significant K-M plots of respective miRNAs (35).

### miRNA target prediction

Gene targets for miRNA were predicted using multiple platforms such as DIANA-microT (36), TargetScan (37), miRDB (38), RNA22 (39). Highest stringency was applied in these tools while predicting the targets and only those targets were considered which were predicted by at least two tools. Further, the expression of these target mRNAs was checked using NCBI (40) GTEx portal (41) and BioGPS (42) to consider only those relevant in endometrial tissue.

## Results

### Differential expression of miRNAs in endometrial cancer (Indian population)

12 control and 32 endometrial cancer samples representing Indian population were subjected to small RNA sequencing and upon differential expression analysis using miRDeep2 algorithm, 927 putative novel miRNAs were identified. Differential expression analysis with a false discovery rate (FDR) <0.001 and 4-fold change (>/=2 log2 fold) in expression, 114 known miRNAs were overexpressed and 81 under expressed in endometrial cancer samples as compared to controls. With similar parameters, 17 novel miRNAs were found to be overexpressed in endometrial cancer.

Table 1 shows a list of differentially expressed miRNAs between control and endometrial cancer samples along with the fold changes in expression.

**Table 1A:** List of Overexpressed miRNAs in Endometrial Cancer in Indian population.

| SI No | miRNA ID | log2FoldChange | p value |
| --- | --- | --- | --- |
| 1 | hsa-miR-10394-3p | 4.054088128 | 2.71E-05 |
| 2 | hsa-miR-10396b-3p | 6.642030444 | 2.78E-08 |
| 3 | hsa-miR-10401-3p | 2.520646231 | 2.16E-08 |
| 4 | hsa-miR-10527-5p | 2.158977733 | 1.09E-10 |
| 5 | hsa-miR-10a-3p | 2.026129502 | 1.48E-05 |
| 6 | hsa-miR-1180-3p | 2.059667981 | 4.98E-08 |
| 7 | hsa-miR-1224-5p | 3.837186398 | 2.54E-08 |
| 8 | hsa-miR-1226-3p | 2.736933859 | 3.49E-06 |
| 9 | hsa-miR-1226-5p | 3.973325539 | 6.51E-07 |
| 10 | hsa-miR-1229-3p | 2.621565248 | 6.09E-05 |
| 11 | hsa-miR-1246 | 6.308484771 | 3E-22 |
| 12 | hsa-miR-1268a | 3.91563877 | 2.78E-14 |
| 13 | hsa-miR-1268b | 3.945837301 | 5.08E-15 |
| 14 | hsa-miR-1269a | 7.555852076 | 2.89E-08 |
| 15 | hsa-miR-1276 | 3.023466992 | 6.54E-06 |
| 16 | hsa-miR-1285-3p | 2.69161633 | 1.68E-14 |
| 17 | hsa-miR-1285-3p | 2.601363856 | 1.54E-13 |
| 18 | hsa-miR-1290 | 4.43946503 | 4.66E-11 |
| 19 | hsa-miR-1293 | 4.907715581 | 2.27E-08 |
| 20 | hsa-miR-1301-3p | 2.093282997 | 4.18E-10 |
| 21 | hsa-miR-1303 | 3.584580146 | 1.86E-22 |
| 22 | hsa-miR-1307-3p | 2.134378342 | 1.19E-16 |
| 23 | hsa-miR-1307-5p | 2.814351367 | 1.01E-12 |
| 24 | hsa-miR-135b-3p | 3.066732372 | 4.12E-07 |
| 25 | hsa-miR-135b-5p | 3.407393464 | 2.19E-11 |
| 26 | hsa-miR-147b-3p | 2.136820314 | 9.9E-06 |
| 27 | hsa-miR-16-2-3p | 2.36429597 | 2.96E-08 |
| 28 | hsa-miR-182-3p | 2.575564427 | 4.43E-06 |
| 29 | hsa-miR-183-3p | 3.445694502 | 9.36E-12 |
| 30 | hsa-miR-183-5p | 3.83824592 | 7.14E-24 |
| 31 | hsa-miR-1908-3p | 3.436181551 | 4.54E-05 |
| 32 | hsa-miR-190b-5p | 2.373047213 | 0.000122 |
| 33 | hsa-miR-200a-5p | 3.050842242 | 3.87E-16 |
| 34 | hsa-miR-200b-3p | 3.33816652 | 2.61E-19 |
| 35 | hsa-miR-200b-5p | 2.518556703 | 1.05E-08 |
| 36 | hsa-miR-200c-5p | 2.364609338 | 2.55E-08 |
| 37 | hsa-miR-205-5p | 4.523590131 | 7.45E-14 |
| 38 | hsa-miR-2276-3p | 2.047247665 | 4.47E-08 |
| 39 | hsa-miR-3074-5p | 2.768100718 | 3.2E-07 |
| 40 | hsa-miR-3127-3p | 2.484067906 | 1.47E-08 |
| 41 | hsa-miR-3176 | 4.644836188 | 8.42E-18 |
| 42 | hsa-miR-3177-3p | 2.068816501 | 4.79E-10 |
| 43 | hsa-miR-3180 | 5.363574971 | 2.34E-10 |
| 44 | hsa-miR-3180-5p | 4.037123627 | 5.39E-07 |
| 45 | hsa-miR-33b-3p | 2.983901364 | 3.8E-05 |
| 46 | hsa-miR-3614-5p | 3.007329912 | 1.28E-08 |
| 47 | hsa-miR-3615 | 3.299140854 | 4.36E-12 |
| 48 | hsa-miR-3620-5p | 3.517970434 | 1.03E-08 |
| 49 | hsa-miR-363-5p | 2.367979953 | 0.000144 |
| 50 | hsa-miR-3651 | 2.12300344 | 2.51E-06 |
| 51 | hsa-miR-3661 | 3.551147644 | 2.44E-05 |
| 52 | hsa-miR-3679-5p | 2.22073617 | 2.19E-07 |
| 53 | hsa-miR-3691-5p | 2.655282781 | 9.54E-07 |
| 54 | hsa-miR-3934-5p | 2.078311438 | 8.89E-11 |
| 55 | hsa-miR-3944-3p | 3.290335189 | 6.14E-11 |
| 56 | hsa-miR-3960 | 5.356861024 | 8.84E-17 |
| 57 | hsa-miR-429 | 3.00457065 | 4.42E-10 |
| 58 | hsa-miR-4301 | 3.82574932 | 9.83E-06 |
| 59 | hsa-miR-4326 | 2.011090912 | 5.8E-05 |
| 60 | hsa-miR-4443 | 2.863216152 | 3.12E-05 |
| 61 | hsa-miR-4446-3p | 3.56757855 | 2E-08 |
| 62 | hsa-miR-4448 | 4.569259283 | 2.94E-12 |
| 63 | hsa-miR-4449 | 2.7865025 | 2.81E-07 |
| 64 | hsa-miR-4474-3p | 2.419310949 | 2.93E-05 |
| 65 | hsa-miR-4485-3p | 3.202173251 | 4.7E-06 |
| 66 | hsa-miR-4488 | 6.241102483 | 7E-22 |
| 67 | hsa-miR-4498 | 3.951245915 | 2.21E-07 |
| 68 | hsa-miR-449a | 3.379747335 | 7.05E-05 |
| 69 | hsa-miR-449c-3p | 5.480856907 | 3.31E-05 |
| 70 | hsa-miR-449c-5p | 3.836057842 | 3.54E-06 |
| 71 | hsa-miR-4508 | 2.810363518 | 1.93E-05 |
| 72 | hsa-miR-4516 | 3.800272979 | 1.1E-06 |
| 73 | hsa-miR-4652-5p | 4.70758974 | 3.61E-09 |
| 74 | hsa-miR-4664-3p | 3.32566877 | 1.05E-14 |
| 75 | hsa-miR-4690-3p | 2.472137501 | 2.93E-05 |
| 76 | hsa-miR-4724-5p | 3.347806657 | 3.57E-08 |
| 77 | hsa-miR-4726-5p | 3.266435935 | 7.25E-05 |
| 78 | hsa-miR-4741 | 2.218386694 | 2.62E-05 |
| 79 | hsa-miR-4746-5p | 3.487769772 | 1.63E-14 |
| 80 | hsa-miR-4758-3p | 2.958652682 | 8.5E-07 |
| 81 | hsa-miR-4767 | 3.587566953 | 7.7E-05 |
| 82 | hsa-miR-4800-3p | 2.652309 | 3.73E-05 |
| 83 | hsa-miR-508-5p | 3.023253696 | 4.52E-06 |
| 84 | hsa-miR-509-5p | 2.604746938 | 3.5E-05 |
| 85 | hsa-miR-514a-3p | 4.871048611 | 2.88E-12 |
| 86 | hsa-miR-522-3p | 3.840460593 | 3.42E-07 |
| 87 | hsa-miR-548ah-3p | 3.350048647 | 7.41E-09 |
| 88 | hsa-miR-548p | 3.319477709 | 9.31E-09 |
| 89 | hsa-miR-5571-3p | 3.901512512 | 4.38E-08 |
| 90 | hsa-miR-5585-3p | 2.773431439 | 2.07E-08 |
| 91 | hsa-miR-592 | 2.485143516 | 4.23E-06 |
| 92 | hsa-miR-615-3p | 5.294008799 | 3.2E-10 |
| 93 | hsa-miR-619-5p | 2.814717272 | 7.07E-09 |
| 94 | hsa-miR-636 | 2.390723152 | 1.42E-05 |
| 95 | hsa-miR-6499-5p | 4.085412696 | 2.99E-05 |
| 96 | hsa-miR-6510-3p | 3.587528103 | 4.1E-08 |
| 97 | hsa-miR-6511b-5p | 2.466528269 | 0.000111 |
| 98 | hsa-miR-6515-5p | 2.020776985 | 9.46E-05 |
| 99 | hsa-miR-6741-3p | 2.634910777 | 4.73E-06 |
| 100 | hsa-miR-6783-3p | 3.218935421 | 3.25E-05 |
| 101 | hsa-miR-6811-5p | 4.056700348 | 0.000125 |
| 102 | hsa-miR-6877-5p | 2.425559917 | 5.26E-07 |
| 103 | hsa-miR-7110-3p | 3.911381814 | 1.73E-05 |
| 104 | hsa-miR-7114-5p | 4.466987024 | 2.77E-05 |
| 105 | hsa-miR-744-5p | 2.050242543 | 1.66E-07 |
| 106 | hsa-miR-760 | 2.457277294 | 1.08E-10 |
| 107 | hsa-miR-7974 | 2.735793233 | 2.41E-06 |
| 108 | hsa-miR-8485 | 3.07401259 | 7.98E-07 |
| 109 | hsa-miR-877-3p | 2.232319229 | 6.23E-07 |
| 110 | hsa-miR-891a-5p | 4.752826748 | 9.75E-07 |
| 111 | hsa-miR-892a | 6.809797754 | 7.6E-09 |
| 112 | hsa-miR-92b-5p | 2.230754457 | 1.15E-07 |
| 113 | hsa-miR-937-3p | 3.547154062 | 3.77E-11 |
| 114 | hsa-miR-940 | 2.672881359 | 1.47E-08 |

**Table 1B:** List of Under-expressed miRNAs in Endometrial Cancer in Indian Population.

| SI No | miRNA ID | log2FoldChange | p value |
| --- | --- | --- | --- |
| 1 | hsa-miR-133b | -4.775595487 | 3.63E-09 |
| 2 | hsa-miR-6507-5p | -4.259067749 | 4.95E-10 |
| 3 | hsa-miR-133a-3p | -3.96258383 | 4.43E-08 |
| 4 | hsa-miR-873-3p | -3.79348598 | 8.74E-08 |
| 5 | hsa-miR-424-5p | -3.781870148 | 1.18E-11 |
| 6 | hsa-miR-99a-5p | -3.774123658 | 1.44E-14 |
| 7 | hsa-miR-542-3p | -3.748479569 | 1.83E-11 |
| 8 | hsa-miR-450b-5p | -3.744000341 | 1.84E-11 |
| 9 | hsa-miR-127-3p | -3.671114999 | 5.54E-19 |
| 10 | hsa-miR-548ba | -3.635903406 | 7.73E-06 |
| 11 | hsa-miR-503-3p | -3.559106178 | 1.09E-07 |
| 12 | hsa-miR-7978 | -3.528104526 | 5.35E-05 |
| 13 | hsa-miR-145-5p | -3.505077547 | 3.7E-12 |
| 14 | hsa-miR-424-3p | -3.379893958 | 4.66E-12 |
| 15 | hsa-miR-376c-3p | -3.347088931 | 1.79E-07 |
| 16 | hsa-miR-1245b-5p | -3.307098492 | 8.47E-08 |
| 17 | hsa-miR-379-5p | -3.295485692 | 1.1E-15 |
| 18 | hsa-miR-542-5p | -3.250572942 | 1.8E-10 |
| 19 | hsa-miR-199b-5p | -3.215391845 | 2.65E-17 |
| 20 | hsa-miR-10b-5p | -3.209943376 | 2.06E-15 |
| 21 | hsa-miR-450a-5p | -3.188018211 | 1.64E-09 |
| 22 | hsa-miR-450a-5p | -3.187895074 | 1.35E-09 |
| 23 | hsa-miR-299-5p | -3.174230526 | 9.95E-13 |
| 24 | hsa-miR-143-5p | -3.172540808 | 4.74E-11 |
| 25 | hsa-miR-2682-5p | -3.103687775 | 0.000132 |
| 26 | hsa-miR-1-3p | -3.005501284 | 1.1E-07 |
| 27 | hsa-miR-1-3p | -3.026403105 | 1.44E-07 |
| 28 | hsa-miR-758-3p | -2.973676546 | 1.69E-13 |
| 29 | hsa-miR-494-3p | -2.968776883 | 1.95E-09 |
| 30 | hsa-miR-450a-2-3p | -2.952995102 | 1.04E-09 |
| 31 | hsa-miR-543 | -2.921320255 | 2.46E-09 |
| 32 | hsa-miR-337-5p | -2.919543949 | 2.69E-08 |
| 33 | hsa-miR-6130 | -2.917549317 | 0.000171 |
| 34 | hsa-miR-379-3p | -2.913258778 | 4.09E-11 |
| 35 | hsa-miR-195-5p | -2.908135863 | 4.1E-14 |
| 36 | hsa-miR-152-3p | -2.853426601 | 3.02E-14 |
| 37 | hsa-miR-1247-3p | -2.838856895 | 2.36E-07 |
| 38 | hsa-miR-668-3p | -2.821040194 | 3.87E-07 |
| 39 | hsa-miR-127-5p | -2.809341694 | 5.35E-09 |
| 40 | hsa-miR-145-3p | -2.763122423 | 1.33E-11 |
| 41 | hsa-miR-431-3p | -2.741124255 | 3.55E-09 |
| 42 | hsa-miR-4510 | -2.740493106 | 2.64E-09 |
| 43 | hsa-miR-889-3p | -2.736550841 | 4.55E-06 |
| 44 | hsa-miR-410-3p | -2.734923047 | 5.07E-07 |
| 45 | hsa-miR-548ag | -2.724428629 | 0.000149 |
| 46 | hsa-miR-337-3p | -2.719261177 | 1.24E-07 |
| 47 | hsa-miR-99a-3p | -2.70206724 | 5.24E-08 |
| 48 | hsa-miR-4636 | -2.694546658 | 1.04E-07 |
| 49 | hsa-miR-381-3p | -2.663998484 | 1.36E-12 |
| 50 | hsa-miR-377-5p | -2.657161529 | 5.12E-09 |
| 51 | hsa-miR-204-5p | -2.650063819 | 6.49E-06 |
| 52 | hsa-miR-487b-3p | -2.64806894 | 3.32E-14 |
| 53 | hsa-miR-493-5p | -2.588245368 | 1.34E-08 |
| 54 | hsa-miR-455-5p | -2.580298559 | 2.34E-11 |
| 55 | hsa-miR-1247-5p | -2.551459633 | 5.5E-07 |
| 56 | hsa-miR-323a-3p | -2.55014504 | 2.17E-09 |
| 57 | hsa-miR-370-5p | -2.547282173 | 1.77E-05 |
| 58 | hsa-let-7c-5p | -2.516242177 | 7.5E-09 |
| 59 | hsa-miR-409-5p | -2.486718091 | 6.35E-09 |
| 60 | hsa-miR-125b-5p | -2.475506379 | 1.44E-09 |
| 61 | hsa-miR-125b-5p | -2.469582719 | 1.71E-09 |
| 62 | hsa-miR-329-3p | -2.455402884 | 1.49E-07 |
| 63 | hsa-miR-136-5p | -2.411387694 | 2.45E-05 |
| 64 | hsa-miR-139-5p | -2.39764053 | 7.86E-12 |
| 65 | hsa-miR-134-5p | -2.34726762 | 2.84E-10 |
| 66 | hsa-miR-140-3p | -2.344745292 | 5.76E-10 |
| 67 | hsa-miR-432-5p | -2.313915209 | 3.48E-07 |
| 68 | hsa-miR-503-5p | -2.30019741 | 2.09E-09 |
| 69 | hsa-miR-382-3p | -2.28995154 | 6.5E-05 |
| 70 | hsa-miR-504-5p | -2.285257202 | 2.79E-06 |
| 71 | hsa-miR-497-5p | -2.281136275 | 3.33E-08 |
| 72 | hsa-miR-143-3p | -2.264085667 | 5.68E-09 |
| 73 | hsa-miR-139-3p | -2.253967178 | 2.81E-05 |
| 74 | hsa-miR-100-5p | -2.23482547 | 1.61E-06 |
| 75 | hsa-miR-199a-3p | -2.187929661 | 4.81E-10 |
| 76 | hsa-miR-199a-3p | -2.187102261 | 4.91E-10 |
| 77 | hsa-miR-199b-3p | -2.187102261 | 4.91E-10 |
| 78 | hsa-miR-675-3p | -2.144157536 | 4.29E-05 |
| 79 | hsa-let-7c-3p | -2.116542091 | 3.55E-05 |
| 80 | hsa-miR-342-3p | -2.114659226 | 7.03E-06 |
| 81 | hsa-miR-214-3p | -2.011042797 | 2.37E-06 |

Table 2 shows a list of novel miRNAs that are differentially expressed in endometrial cancer.

**Table 2:**
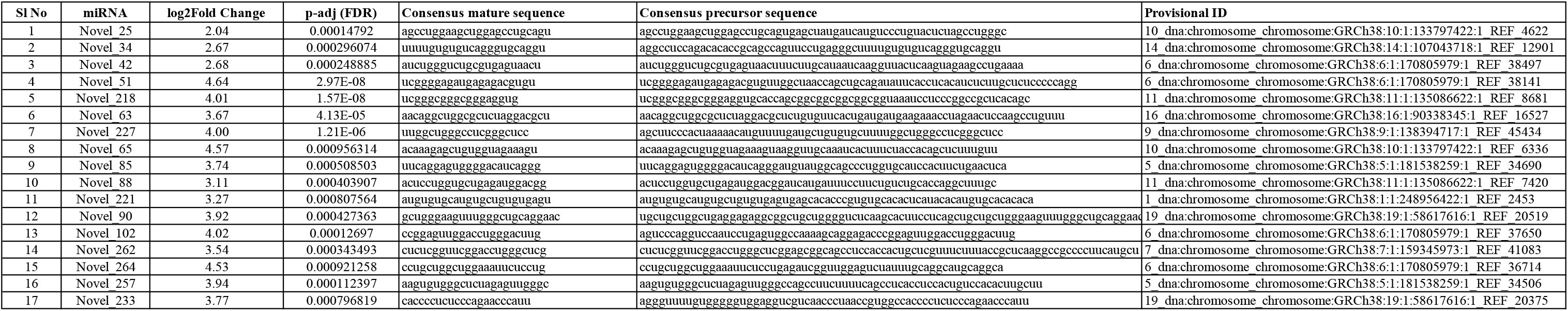
List of differentially expressed novel miRNAs in endometrial cancer.

### Comparison with TCGA data

In order to understand the relevance of the differentially expressed miRNAs in prognosis, the list of differentially expressed miRNAs in Indian population was compared to the differentially expressed miRNA data from TCGA. Among the two endometrial cancer data sets available from TCGA, the one which had information on miRNA expression as well as clinical follow up was chosen (dataset ID: TCGA.UCEC.sampleMap/miRNA_HiSeq_gene). The TCGA endometrioid cancer cohort comprises of data collected from 429 patient samples using the IlluminaHiseq platform. Out of these, 398 were tumor samples, and 31 were control samples. Clinical information included sample type, histological type, histological grade, age and year of initial pathological diagnosis, height, weight, clinical stage, pregnancy status, menopause status, birth control pill usage history, diabetes, additional treatment completion success outcome, vital status, etc. 2238 miRNAs were found to be differentially expressed between control and endometrial cancer according to the TCGA data. When compared to data from our cohort, 91 overexpressed and 77 under-expressed miRNAs were also found in the TCGA data (Figure 1 and Table 3). However, in the common list, 35 miRNAs were overexpressed in both data, 63 miRNAs were under-expressed in both data. For the remaining miRNAs, the fold changes did not match between our data and TCGA data.

**Table 3A:** List of common miRNAs Overexpressed in Endometrial Cancer.

| Sl No | miRNA Accession ID | miRNA ID |
| --- | --- | --- |
| 1 | MIMAT0004555 | hsa-miR-10a-3p |
| 2 | MIMAT0005825 | hsa-miR-1180-3p |
| 3 | MIMAT0005458 | hsa-miR-1224-5p |
| 4 | MIMAT0005577 | hsa-miR-1226-3p |
| 5 | MIMAT0005584 | hsa-miR-1229-3p |
| 6 | MIMAT0005898 | hsa-miR-1246 |
| 7 | MIMAT0018925 | hsa-miR-1268b |
| 8 | MIMAT0005923 | hsa-miR-1269a |
| 9 | MIMAT0005930 | hsa-miR-1276 |
| 10 | MIMAT0005876 | hsa-miR-1285-3p |
| 11 | MIMAT0005797 | hsa-miR-1301-3p |
| 12 | MIMAT0005951 | hsa-miR-1307-3p |
| 13 | MIMAT0022727 | hsa-miR-1307-5p |
| 14 | MIMAT0004698 | hsa-miR-135b-3p |
| 15 | MIMAT0000758 | hsa-miR-135b-5p |
| 16 | MIMAT0004928 | hsa-miR-147b-3p |
| 17 | MIMAT0004518 | hsa-miR-16-2-3p |
| 18 | MIMAT0000260 | hsa-miR-182-3p |
| 19 | MIMAT0004560 | hsa-miR-183-3p |
| 20 | MIMAT0000261 | hsa-miR-183-5p |
| 21 | MIMAT0026916 | hsa-miR-1908-3p |
| 22 | MIMAT0004929 | hsa-miR-190b-5p |
| 23 | MIMAT0001620 | hsa-miR-200a-5p |
| 24 | MIMAT0000318 | hsa-miR-200b-3p |
| 25 | MIMAT0004571 | hsa-miR-200b-5p |
| 26 | MIMAT0004657 | hsa-miR-200c-5p |
| 27 | MIMAT0000266 | hsa-miR-205-5p |
| 28 | MIMAT0011775 | hsa-miR-2276-3p |
| 29 | MIMAT0019208 | hsa-miR-3074-5p |
| 30 | MIMAT0019201 | hsa-miR-3127-3p |
| 31 | MIMAT0015053 | hsa-miR-3176 |
| 32 | MIMAT0015054 | hsa-miR-3177-3p |
| 33 | MIMAT0018178 | hsa-miR-3180 |
| 34 | MIMAT0004811 | hsa-miR-33b-3p |
| 35 | MIMAT0017992 | hsa-miR-3614-5p |
| 36 | MIMAT0017994 | hsa-miR-3615 |
| 37 | MIMAT0022967 | hsa-miR-3620-5p |
| 38 | MIMAT0003385 | hsa-miR-363-5p |
| 39 | MIMAT0018071 | hsa-miR-3651 |
| 40 | MIMAT0018082 | hsa-miR-3661 |
| 41 | MIMAT0018104 | hsa-miR-3679-5p |
| 42 | MIMAT0018120 | hsa-miR-3691-5p |
| 43 | MIMAT0018349 | hsa-miR-3934-5p |
| 44 | MIMAT0018360 | hsa-miR-3944-3p |
| 45 | MIMAT0001536 | hsa-miR-429 |
| 46 | MIMAT0016888 | hsa-miR-4326 |
| 47 | MIMAT0018961 | hsa-miR-4443 |
| 48 | MIMAT0018965 | hsa-miR-4446-3p |
| 49 | MIMAT0018968 | hsa-miR-4449 |
| 50 | MIMAT0019022 | hsa-miR-4488 |
| 51 | MIMAT0019033 | hsa-miR-4498 |
| 52 | MIMAT0001541 | hsa-miR-449a |
| 53 | MIMAT0013771 | hsa-miR-449c-3p |
| 54 | MIMAT0010251 | hsa-miR-449c-5p |
| 55 | MIMAT0019045 | hsa-miR-4508 |
| 56 | MIMAT0019053 | hsa-miR-4516 |
| 57 | MIMAT0019716 | hsa-miR-4652-5p |
| 58 | MIMAT0019738 | hsa-miR-4664-3p |
| 59 | MIMAT0019780 | hsa-miR-4690-3p |
| 60 | MIMAT0019841 | hsa-miR-4724-5p |
| 61 | MIMAT0019845 | hsa-miR-4726-5p |
| 62 | MIMAT0019871 | hsa-miR-4741 |
| 63 | MIMAT0019880 | hsa-miR-4746-5p |
| 64 | MIMAT0019904 | hsa-miR-4758-3p |
| 65 | MIMAT0019919 | hsa-miR-4767 |
| 66 | MIMAT0019979 | hsa-miR-4800-3p |
| 67 | MIMAT0004778 | hsa-miR-508-5p |
| 68 | MIMAT0004779 | hsa-miR-509-5p |
| 69 | MIMAT0002883 | hsa-miR-514a-3p |
| 70 | MIMAT0002868 | hsa-miR-522-3p |
| 71 | MIMAT0022258 | hsa-miR-5571-3p |
| 72 | MIMAT0003260 | hsa-miR-592 |
| 73 | MIMAT0003283 | hsa-miR-615-3p |
| 74 | MIMAT0003306 | hsa-miR-636 |
| 75 | MIMAT0025450 | hsa-miR-6499-5p |
| 76 | MIMAT0025477 | hsa-miR-6510-3p |
| 77 | MIMAT0025486 | hsa-miR-6515-5p |
| 78 | MIMAT0027384 | hsa-miR-6741-3p |
| 79 | MIMAT0027467 | hsa-miR-6783-3p |
| 80 | MIMAT0027654 | hsa-miR-6877-5p |
| 81 | MIMAT0028118 | hsa-miR-7110-3p |
| 82 | MIMAT0028125 | hsa-miR-7114-5p |
| 83 | MIMAT0004945 | hsa-miR-744-5p |
| 84 | MIMAT0004957 | hsa-miR-760 |
| 85 | MIMAT0031177 | hsa-miR-7974 |
| 86 | MIMAT0004950 | hsa-miR-877-3p |
| 87 | MIMAT0004902 | hsa-miR-891a-5p |
| 88 | MIMAT0004907 | hsa-miR-892a |
| 89 | MIMAT0004792 | hsa-miR-92b-5p |
| 90 | MIMAT0004980 | hsa-miR-937-3p |
| 91 | MIMAT0004983 | hsa-miR-940 |

**Figure 1:**
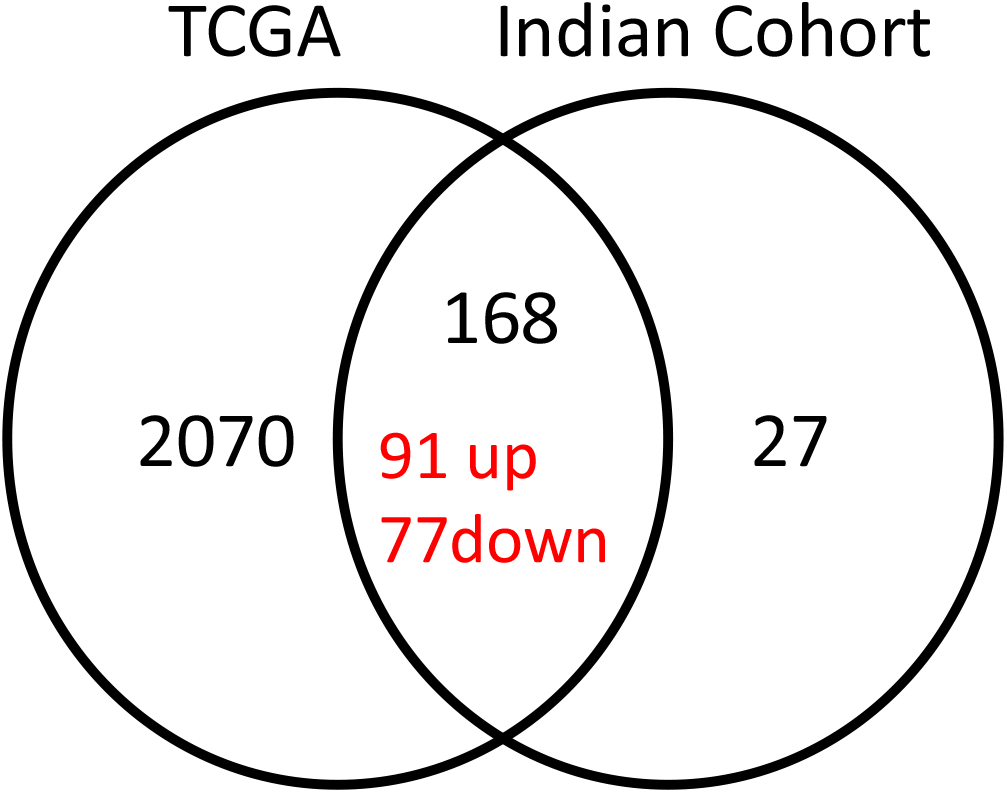
Differentially expressed miRNAs between control and cancerous endometrial tissue in TCGA data set and Indian cohort. 168 miRNAs are found in common to both data sets. 2070 are exclusive to TCGA and 27 are exclusive to Indian patients.

### Survival Analysis

The survival analysis of common miRNAs was performed as described in the methods. This revealed 40 overexpressed miRNAs (out of 91) and 25 under-expressed miRNAs (out of 77) had a significant effect on survival. 20 miRNAs show better survival when overexpressed in tumors (Figure 2A, Table 4A), and 45 show poorer survival when overexpressed in tumors (Figure 2B, Table 4B).

**Table 3B:** List of common miRNAs Under-expressed in Endometrial Cancer.

| SI No | miRNA Accession ID | miRNA ID |
| --- | --- | --- |
| 1 | MIMAT0002890 | hsa-miR-299-5p |
| 2 | MIMAT0019047 | hsa-miR-4510 |
| 3 | MIMAT0019693 | hsa-miR-4636 |
| 4 | MIMAT0024614 | hsa-miR-6130 |
| 5 | MIMAT0000448 | hsa-miR-136-5p |
| 6 | MIMAT0000446 | hsa-miR-127-3p |
| 7 | MIMAT0000447 | hsa-miR-134-5p |
| 8 | MIMAT0026472 | hsa-let-7c-3p |
| 9 | MIMAT0001545 | hsa-miR-450a-5p |
| 10 | MIMAT0022721 | hsa-miR-1247-3p |
| 11 | MIMAT0003150 | hsa-miR-455-5p |
| 12 | MIMAT0002171 | hsa-miR-410-3p |
| 13 | MIMAT0000736 | hsa-miR-381-3p |
| 14 | MIMAT0000733 | hsa-miR-379-5p |
| 15 | MIMAT0002820 | hsa-miR-497-5p |
| 16 | MIMAT0000461 | hsa-miR-195-5p |
| 17 | MIMAT0031181 | hsa-miR-7978 |
| 18 | MIMAT0003389 | hsa-miR-542-3p |
| 19 | MIMAT0031074 | hsa-miR-450a-2-3p |
| 20 | MIMAT0000720 | hsa-miR-376c-3p |
| 21 | MIMAT0003881 | hsa-miR-668-3p |
| 22 | MIMAT0004604 | hsa-miR-127-5p |
| 23 | MIMAT0004601 | hsa-miR-145-3p |
| 24 | MIMAT0004511 | hsa-miR-99a-3p |
| 25 | MIMAT0000265 | hsa-miR-204-5p |
| 26 | MIMAT0000263 | hsa-miR-199b-5p |
| 27 | MIMAT0000064 | hsa-let-7c-5p |
| 28 | MIMAT0003180 | hsa-miR-487b-3p |
| 29 | MIMAT0003340 | hsa-miR-542-5p |
| 30 | MIMAT0019950 | hsa-miR-1245b-5p |
| 31 | MIMAT0000271 | hsa-miR-214-3p |
| 32 | MIMAT0002814 | hsa-miR-432-5p |
| 33 | MIMAT0002816 | hsa-miR-494-3p |
| 34 | MIMAT0002813 | hsa-miR-493-5p |
| 35 | MIMAT0000416 | hsa-miR-1-3p |
| 36 | MIMAT0000098 | hsa-miR-100-5p |
| 37 | MIMAT0000097 | hsa-miR-99a-5p |
| 38 | MIMAT0004757 | hsa-miR-431-3p |
| 39 | MIMAT0018969 | hsa-miR-548ag |
| 40 | MIMAT0025470 | hsa-miR-6507-5p |
| 41 | MIMAT0004749 | hsa-miR-424-3p |
| 42 | MIMAT0000427 | hsa-miR-133a-3p |
| 43 | MIMAT0000423 | hsa-miR-125b-5p |
| 44 | MIMAT0004563 | hsa-miR-199b-3p |
| 45 | MIMAT0022697 | hsa-miR-382-3p |
| 46 | MIMAT0002875 | hsa-miR-504-5p |
| 47 | MIMAT0002874 | hsa-miR-503-5p |
| 48 | MIMAT0000254 | hsa-miR-10b-5p |
| 49 | MIMAT0000250 | hsa-miR-139-5p |
| 50 | MIMAT0000438 | hsa-miR-152-3p |
| 51 | MIMAT0000435 | hsa-miR-143-3p |
| 52 | MIMAT0000437 | hsa-miR-145-5p |
| 53 | MIMAT0004552 | hsa-miR-139-3p |
| 54 | MIMAT0004954 | hsa-miR-543 |
| 55 | MIMAT0022925 | hsa-miR-503-3p |
| 56 | MIMAT0013517 | hsa-miR-2682-5p |
| 57 | MIMAT0006790 | hsa-miR-675-3p |
| 58 | MIMAT0001638 | hsa-miR-409-5p |
| 59 | MIMAT0000232 | hsa-miR-199a-3p |
| 60 | MIMAT0003879 | hsa-miR-758-3p |
| 61 | MIMAT0026483 | hsa-miR-370-5p |
| 62 | MIMAT0004690 | hsa-miR-379-3p |
| 63 | MIMAT0031175 | hsa-miR-548ba |
| 64 | MIMAT0001629 | hsa-miR-329-3p |
| 65 | MIMAT0005899 | hsa-miR-1247-5p |
| 66 | MIMAT0000754 | hsa-miR-337-3p |
| 67 | MIMAT0000755 | hsa-miR-323a-3p |
| 68 | MIMAT0000753 | hsa-miR-342-3p |
| 69 | MIMAT0004921 | hsa-miR-889-3p |
| 70 | MIMAT0001341 | hsa-miR-424-5p |
| 71 | MIMAT0004597 | hsa-miR-140-3p |
| 72 | MIMAT0004599 | hsa-miR-143-5p |
| 73 | MIMAT0004689 | hsa-miR-377-5p |
| 74 | MIMAT0000770 | hsa-miR-133b |
| 75 | MIMAT0004909 | hsa-miR-450b-5p |
| 76 | MIMAT0004695 | hsa-miR-337-5p |
| 77 | MIMAT0022717 | hsa-miR-873-3p |

**Table 4A:** List of miRNAs which show positive correlation with Overall survival (Higher expression, better survival)

| Sl No | miRNA Accession ID | miRNA ID | log2FoldChange (TCGA) | Hazard Ratio | Class Interval |
| --- | --- | --- | --- | --- | --- |
| 1 | MIMAT0005825 | hsa-miR-1180-3p | 1.676071744 | 0.56 | 0.34-0.92 |
| 2 | MIMAT0005458 | hsa-miR-1224-5p | 1.606135527 | 0.49 | 0.3-0.79 |
| 3 | MIMAT0004929 | hsa-miR-190b-5p | 0.733345776 | 0.38 | 0.22-0.64 |
| 4 | MIMAT0001620 | hsa-miR-200a-5p | 4.730269869 | 0.53 | 0.32-0.86 |
| 5 | MIMAT0004571 | hsa-miR-200b-5p | 3.799513654 | 0.51 | 0.31-0.85 |
| 6 | MIMAT0019208 | hsa-miR-3074-5p | 2.515021136 | 0.57 | 0.35-0.93 |
| 7 | MIMAT0015054 | hsa-miR-3177-3p | 0.328884247 | 0.58 | 0.35-0.96 |
| 8 | MIMAT0004811 | hsa-miR-33b-3p | 0.517516099 | 0.58 | 0.36-0.94 |
| 9 | MIMAT0017992 | hsa-miR-3614-5p | 1.814518867 | 0.25 | 0.11-0.57 |
| 10 | MIMAT0001541 | hsa-miR-449a | 2.602895724 | 0.5 | 0.32-0.86 |
| 11 | MIMAT0010251 | hsa-miR-449c-5p | 2.067851364 | 0.55 | 0.32-0.95 |
| 12 | MIMAT0019880 | hsa-miR-4746-5p | 2.028831966 | 0.45 | 0.28-0.72 |
| 13 | MIMAT0004945 | hsa-miR-744-5p | 1.705368231 | 0.59 | 0.36-0.95 |
| 14 | MIMAT0022721 | hsa-miR-1247-3p | -2.957314443 | 0.45 | 0.24-0.82 |
| 15 | MIMAT0005899 | hsa-miR-1247-5p | -1.826285517 | 0.6 | 0.37-0.97 |
| 16 | MIMAT0004597 | hsa-miR-140-3p | -1.415767178 | 0.46 | 0.232-0.89 |
| 17 | MIMAT0004601 | hsa-miR-145-3p | -2.515285219 | 0.6 | 0.37-0.98 |
| 18 | MIMAT0000437 | hsa-miR-145-5p | -2.569189935 | 0.56 | 0.34-0.93 |
| 19 | MIMAT0000461 | hsa-miR-195-5p | -1.708796623 | 0.58 | 0.35-0.95 |
| 20 | MIMAT0002820 | hsa-miR-497-5p | -0.846226612 | 0.49 | 0.3-0.81 |

**Table 4B:** List of miRNAs which show negative correlation with Overall survival (Higher expression, poor survival)

| **Table 4B: List of miRNAs which show negative correlation with Overall survival** |  |  |  |  |  |
| --- | --- | --- | --- | --- | --- |
| **(Higher expression, poor survival)** |  |  |  |  |  |
| **Sl No** | **miRNA Accession ID** | **miRNA ID** | **log2FoldChange (TCGA)** | **Hazard Ratio** | **Class Interval** |
| 1 | MIMAT0004555 | hsa-miR-10a-3p | 1.942551751 | 1.79 | 1.11-2.88 |
| 2 | MIMAT0005577 | hsa-miR-1226-3p | 1.325461708 | 2.02 | 1.24-3.29 |
| 3 | MIMAT0005584 | hsa-miR-1229-3p | 0.770005091 | 2.09 | 1.3-3.35 |
| 4 | MIMAT0005923 | hsa-miR-1269a | 3.156363022 | 2.17 | 1.35-3.49 |
| 5 | MIMAT0005930 | hsa-miR-1276 | 0.512369898 | 2.42 | 1.35-4.37 |
| 6 | MIMAT0005797 | hsa-miR-1301-3p | 2.656312442 | 2.87 | 1.47-5.63 |
| 7 | MIMAT0004518 | hsa-miR-16-2-3p | 1.905243879 | 1.77 | 1.05-2.98 |
| 8 | MIMAT0000260 | hsa-miR-182-3p | 0.503988851 | 2.18 | 1.22-3.89 |
| 9 | MIMAT0019201 | hsa-miR-3127-3p | 0.431996383 | 1.69 | 1.03-2.76 |
| 10 | MIMAT0015053 | hsa-miR-3176 | 0.43234325 | 1.71 | 1.06-2.75 |
| 11 | MIMAT0018120 | hsa-miR-3691-5p | 0.604429657 | 2.33 | 1.19-4.55 |
| 12 | MIMAT0019780 | hsa-miR-4690-3p | 0.201513485 | 1.81 | 1.13-2.91 |
| 13 | MIMAT0019904 | hsa-miR-4758-3p | 0.550324869 | 1.96 | 1.22-3.15 |
| 14 | MIMAT0002883 | hsa-miR-514a-3p | 0.42044708 | 2.29 | 1.4-3.77 |
| 15 | MIMAT0002868 | hsa-miR-522-3p | 0.93814227 | 1.96 | 1.21-3.17 |
| 16 | MIMAT0003260 | hsa-miR-592 | 2.209988792 | 1.71 | 1.04-2.81 |
| 17 | MIMAT0025450 | hsa-miR-6499-5p | 0.685171007 | 1.94 | 1.15-3.26 |
| 18 | MIMAT0027467 | hsa-miR-6783-3p | 0.313200902 | 1.83 | 1.12-2.98 |
| 19 | MIMAT0028118 | hsa-miR-7110-3p | 0.271280283 | 1.82 | 1.02-3.22 |
| 20 | MIMAT0028125 | hsa-miR-7114-5p | 0.295414768 | 1.89 | 1.17-3.07 |
| 21 | MIMAT0004957 | hsa-miR-760 | 1.226040877 | 1.99 | 1.22-3.23 |
| 22 | MIMAT0031177 | hsa-miR-7974 | 0.620317931 | 1.97 | 1.22-3.17 |
| 23 | MIMAT0004950 | hsa-miR-877-3p | 0.547782614 | 2.01 | 1.06-3.84 |
| 24 | MIMAT0004902 | hsa-miR-891a-5p | 2.417101688 | 1.72 | 1.07-2.76 |
| 25 | MIMAT0004907 | hsa-miR-892a | 0.942158295 | 1.6 | 1-2.58 |
| 26 | MIMAT0004980 | hsa-miR-937-3p | 1.623371751 | 2.12 | 1.26-3.52 |
| 27 | MIMAT0004983 | hsa-miR-940 | 2.154979634 | 1.78 | 1.1-2.87 |
| 28 | MIMAT0000423 | hsa-miR-125b-5p | -1.444726998 | 1.71 | 1.05-2.78 |
| 29 | MIMAT0000755 | hsa-miR-323a-3p | 0.306883982 | 1.85 | 1.15-2.98 |
| 30 | MIMAT0001629 | hsa-miR-329-3p | -0.057402947 | 1.92 | 1.22-3.09 |
| 31 | MIMAT0026483 | hsa-miR-370-5p | -0.590254896 | 1.82 | 1.12-2.93 |
| 32 | MIMAT0004690 | hsa-miR-379-3p | -0.059253436 | 1.67 | 1-2.79 |
| 33 | MIMAT0000736 | hsa-miR-381-3p | -2.064472846 | 1.7 | 1.06-2.74 |
| 34 | MIMAT0002171 | hsa-miR-410-3p | -0.699074669 | 1.87 | 1.12-3.13 |
| 35 | MIMAT0004757 | hsa-miR-431-3p | -0.218727339 | 1.85 | 1.08-3.17 |
| 36 | MIMAT0019047 | hsa-miR-4510 | -0.051697652 | 1.77 | 1.05-3.01 |
| 37 | MIMAT0019693 | hsa-miR-4636 | 0.148741073 | 2.11 | 1.15-3.87 |
| 38 | MIMAT0003180 | hsa-miR-487b-3p | -0.469981718 | 1.77 | 1.07-2.92 |
| 39 | MIMAT0002813 | hsa-miR-493-5p | 0.01375935 | 1.89 | 1.03-3.46 |
| 40 | MIMAT0002875 | hsa-miR-504-5p | 0.288675864 | 2.33 | 1.45-3.74 |
| 41 | MIMAT0003340 | hsa-miR-542-5p | -1.48100984 | 1.62 | 1-2.61 |
| 42 | MIMAT0031175 | hsa-miR-548ba | 0.015311909 | 1.77 | 1.1-2.86 |
| 43 | MIMAT0025470 | hsa-miR-6507-5p | -0.121791708 | 1.61 | 1-2.58 |
| 44 | MIMAT0022717 | hsa-miR-873-3p | 0.16217544 | 1.83 | 1.13-2.94 |
| 45 | MIMAT0004511 | hsa-miR-99a-3p | -1.005905322 | 1.99 | 1.12-3.54 |

**Figure 2:**
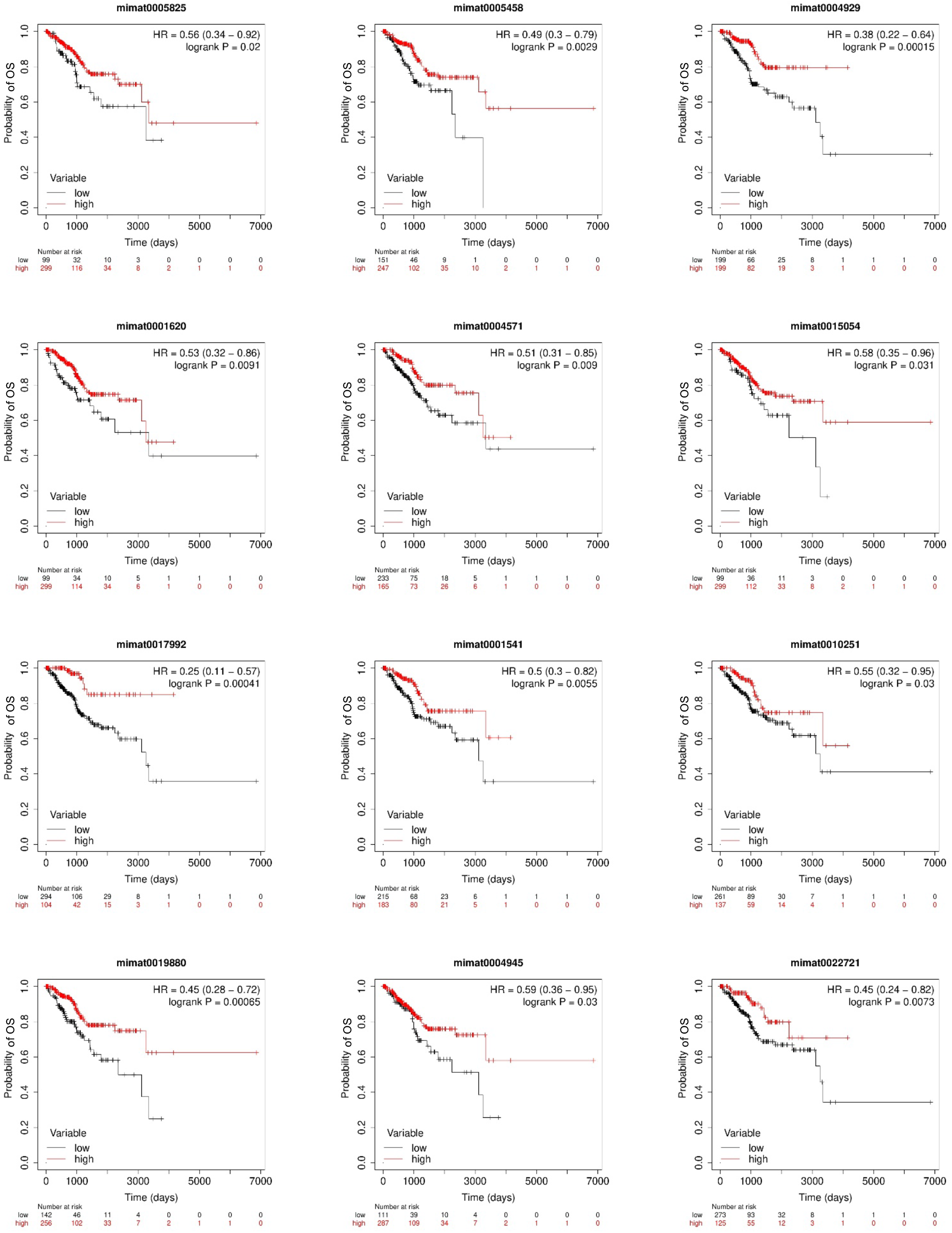
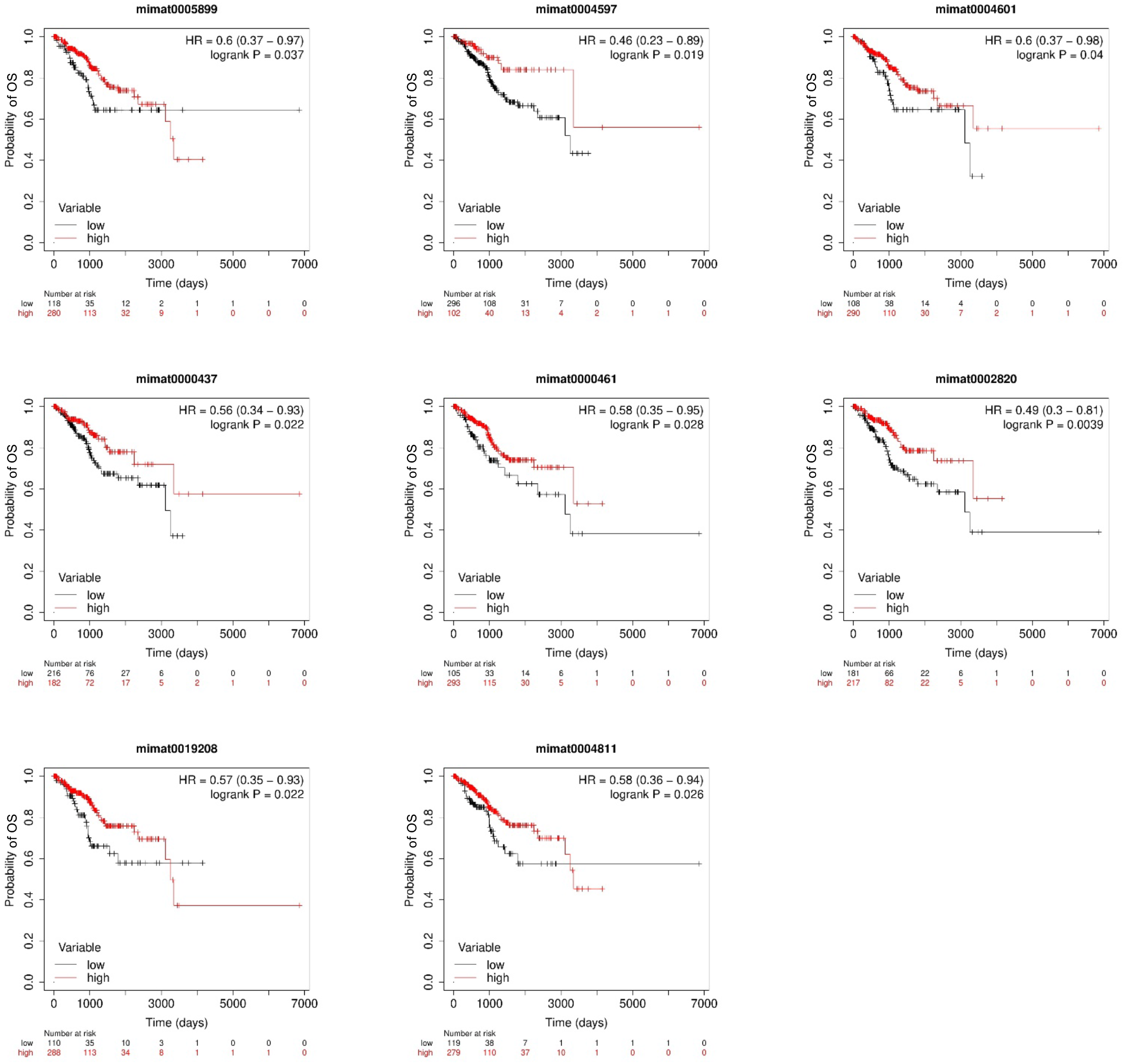
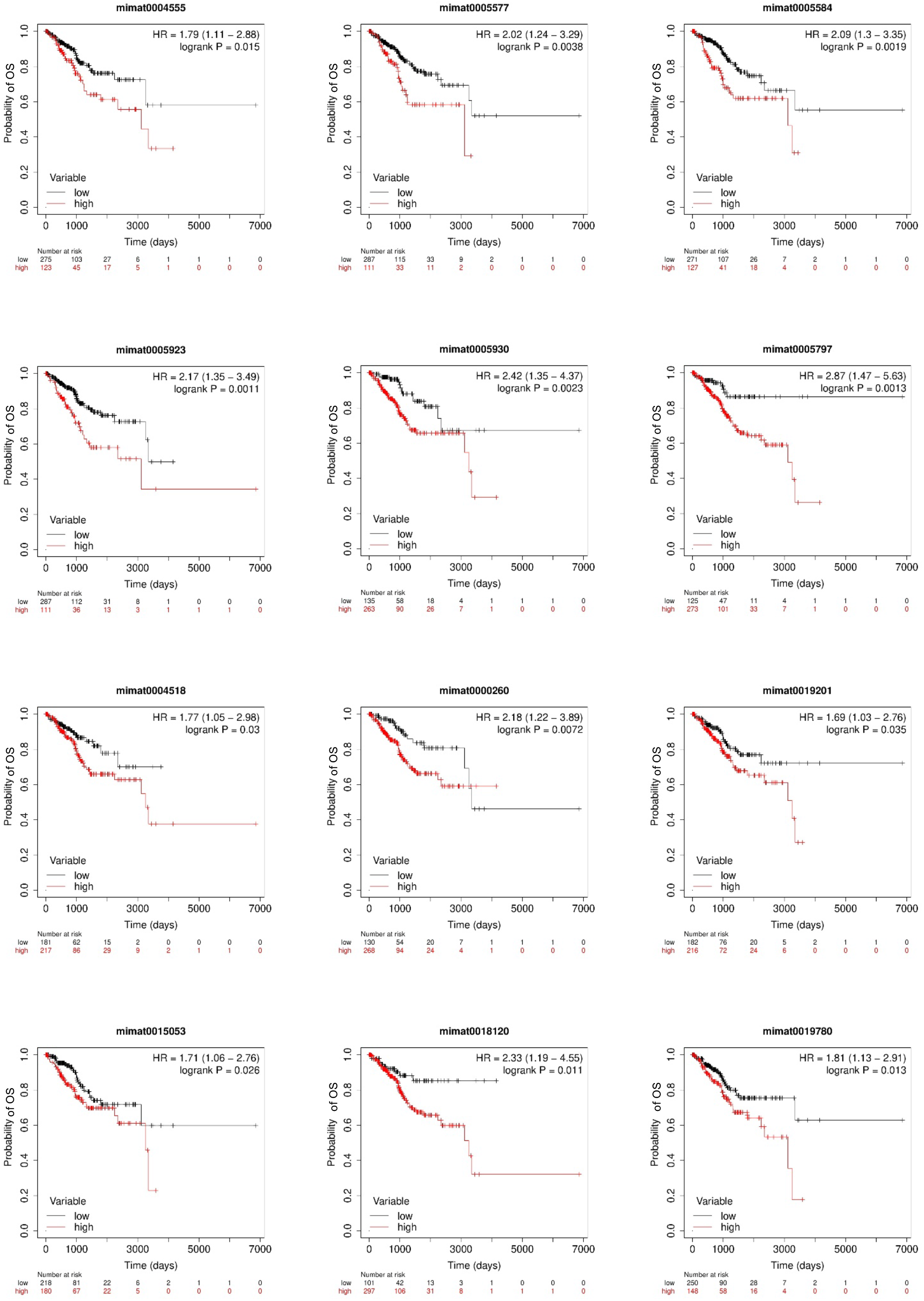
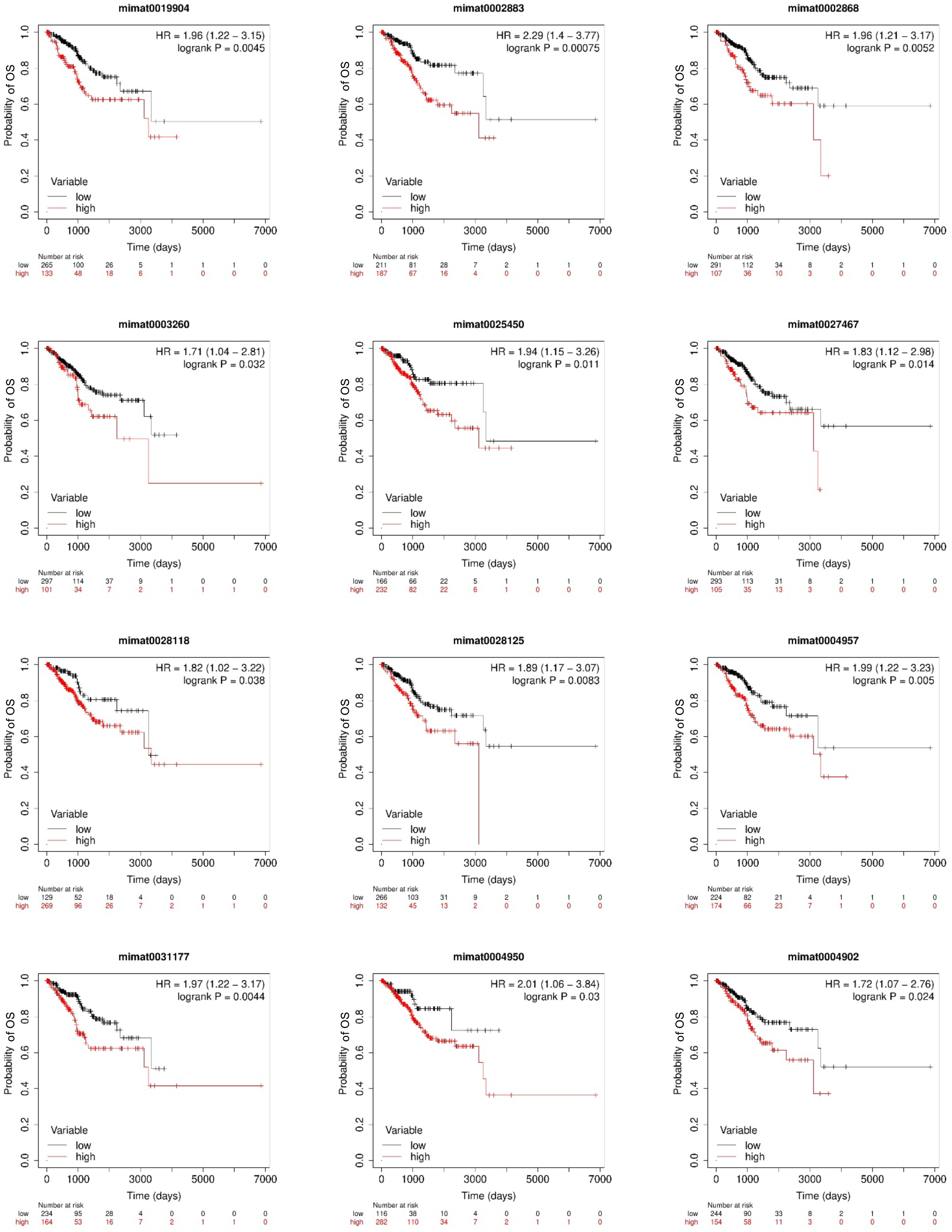
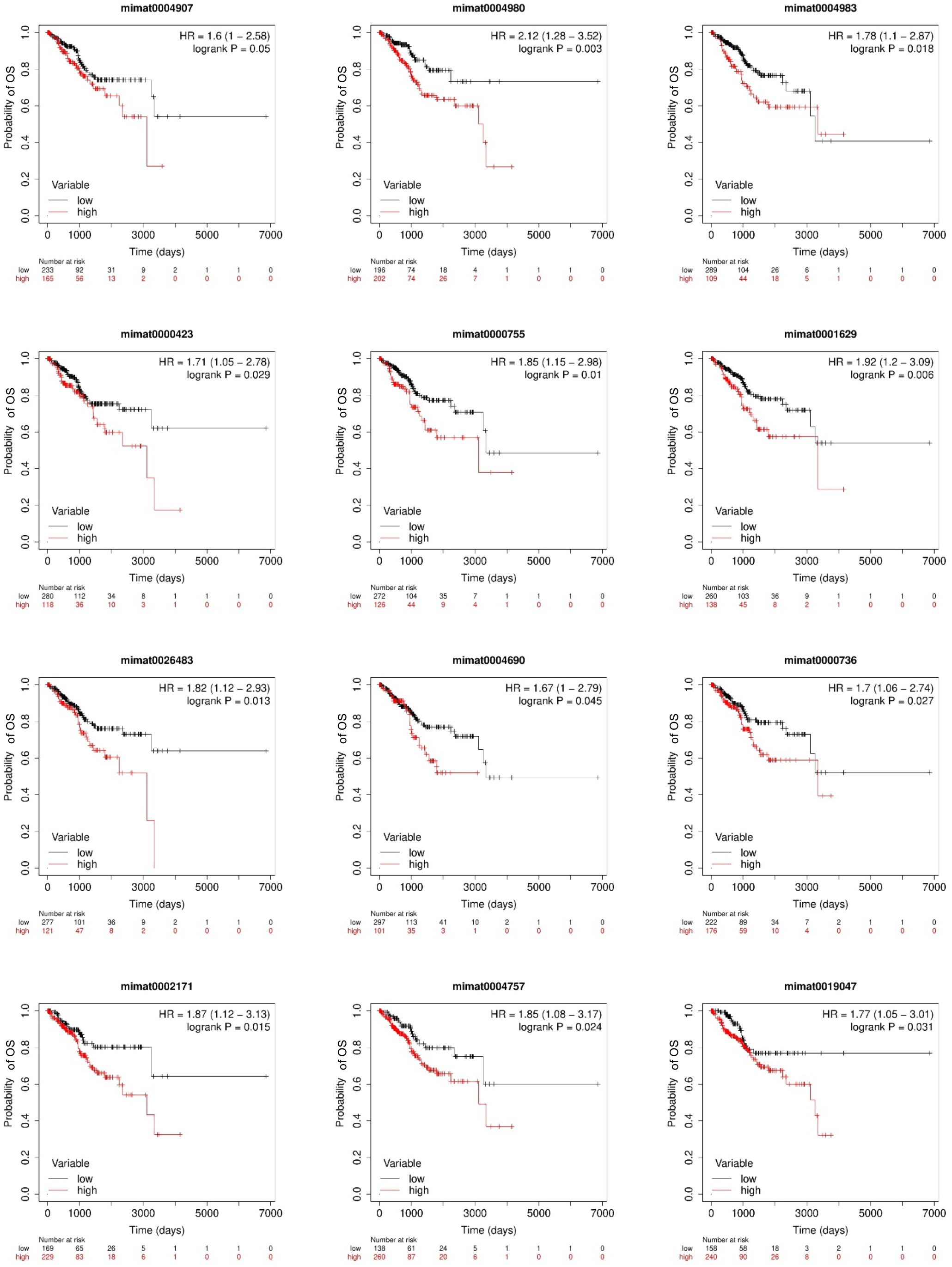
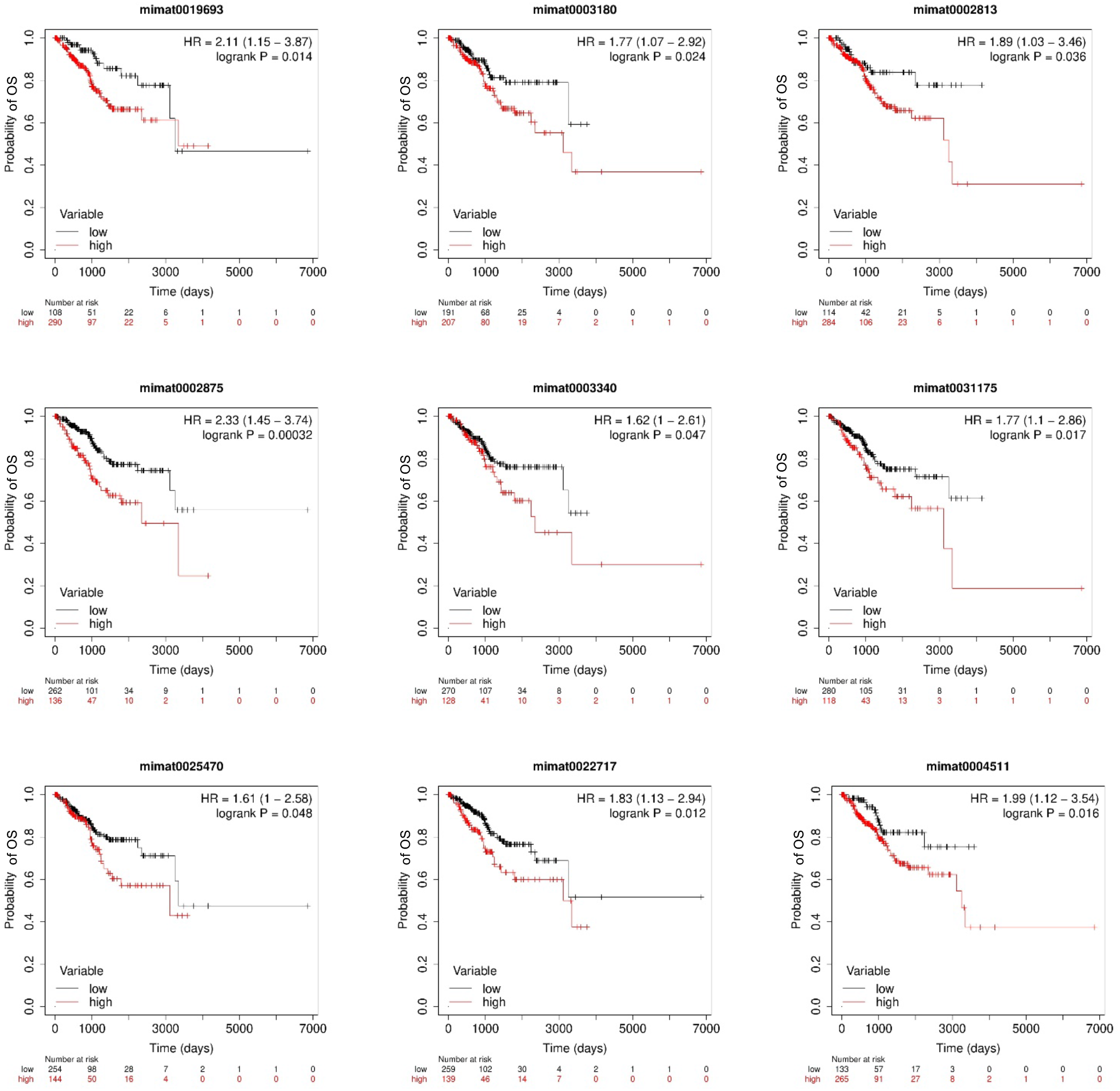
Kaplan Meier Plots of miRNAs which have an impact on overall survival. 2A. K-M plots of miRNAs which are positively correlated with survival (higher expression correlates with better survival) 2B. K-M plots of miRNAs which are negatively correlated with survival (higher expression correlates with poor survival)

### Prediction of targets of the miRNAs

mRNA targets were predicted for the miRNAs which had significant impact on survival. These targets have been predicted using multiple tools such DIANA-microT, TargetScan and miRDB, RNA22 as described in methods (Figure 3). Targets which were predicted by at least two tools were considered for further analysis. The mRNA targets were further filtered using information from NCBI, GTEx and BioGPS for its expression in endometrium. After these two filters we found 40 mRNA targets for overexpressed miRNAs (Table 5A) and 25 mRNA targets for under-expressed miRNAs (Table 5B). The mRNA targets were then compared to TCGA data set for their status in endometrial cancer. From this, it was found that 59 mRNA (targets of 34 overexpressed and 25 under-expressed miRNAs) targets were present in TCGA data. We then analyzed the miRNA-mRNA pairs and found that a total 16 pairs of miRNA-mRNA showed inverse correlation, which could be potential biomarkers for endometrial cancer prognosis (Table 6).

**Figure 3:**
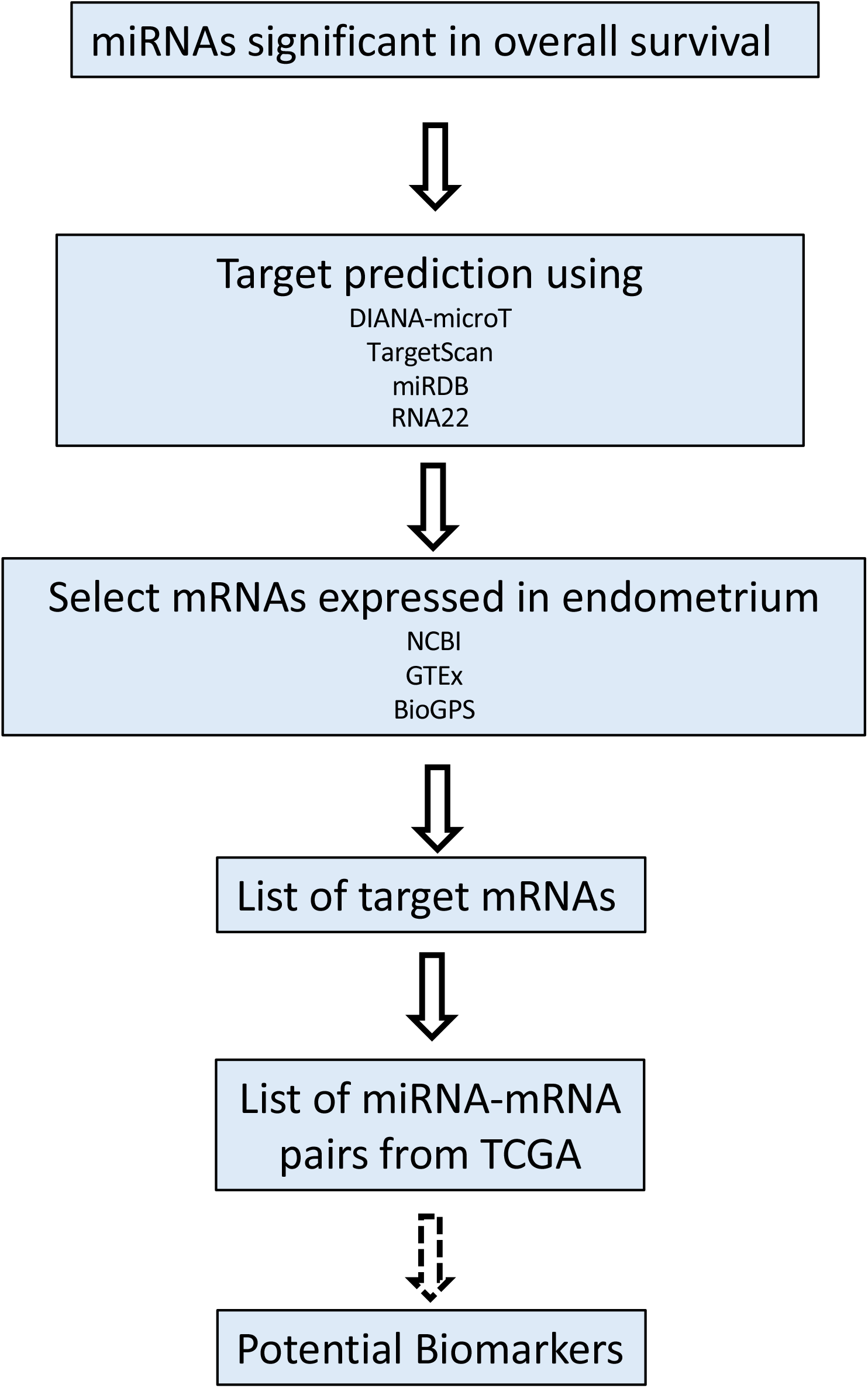
Schematic work flow for identification of mRNA targets of the miRNAs

**Table 5A:** List of predicted mRNA targets for the Overexpressed miRNAs which have significant impact on Overall survival.

| SI No | miRNA Accession ID | miRNA ID | Target Gene |  |
| --- | --- | --- | --- | --- |
| 1 | MIMAT0004555 | hsa-miR-10a-3p | PTEN | Phosphatase and tensin homolog |
| 2 | MIMAT0005825 | hsa-miR-1180-3p | BCL6 | B-cell lymphoma 6 protein |
| 3 | MIMAT0005458 | hsa-miR-1224-5p | SIRT6 | Sirtuin 6 |
| 4 | MIMAT0005577 | hsa-miR-1226-3p | NRAS | Neuroblastoma RAS viral oncogene homolog |
| 5 | MIMAT0005584 | hsa-miR-1229-3p | CCNT1 | Cyclin T1 |
| 6 | MIMAT0005923 | hsa-miR-1269a | KMT2A | Lysine methyltransferase 2A (MLL) |
| 7 | MIMAT0005930 | hsa-miR-1276 | DDX5 | DEAD-box helicase 5 (p68) |
| 8 | MIMAT0005797 | hsa-miR-1301-3p | ANK2 | Ankyrin 2 |
| 9 | MIMAT0004518 | hsa-miR-16-2-3p | CDK6 | Cyclin-dependent kinase 6 |
| 10 | MIMAT0000260 | hsa-miR-182-3p | RUNX1T | RUNX1 partner transcriptional co-repressor 1 |
| 11 | MIMAT0004929 | hsa-miR-190b-5p | TNRC6B | Trinucleotide repeat-containing gene 6B protein |
| 12 | MIMAT0001620 | hsa-miR-200a-5p | ZEB2 | Zinc finger E-box-binding homeobox 2 (SIP1) |
| 13 | MIMAT0004571 | hsa-miR-200b-5p | CKS2 | Cyclin-dependent kinases regulatory subunit 2 |
| 14 | MIMAT0019208 | hsa-miR-3074-5p | THBS1 | Thrombospondin 1 |
| 15 | MIMAT0019201 | hsa-miR-3127-3p | WNT5A | Wingless-type MMTV integration site family 5A |
| 16 | MIMAT0015053 | hsa-miR-3176 | BCL6 | B-cell lymphoma 6 protein |
| 17 | MIMAT0015054 | hsa-miR-3177-3p | PRKCG | Protein kinase C gamma |
| 18 | MIMAT0004811 | hsa-miR-33b-3p | PTGER3 | Prostaglandin E receptor 3 (subtype EP3) |
| 19 | MIMAT0017992 | hsa-miR-3614-5p | MLH1 | MutL homolog 1 |
| 20 | MIMAT0018120 | hsa-miR-3691-5p | RASGRP2 | RAS guanyl-releasing protein 2 |
| 21 | MIMAT0001541 | hsa-miR-449a | LEF1 | Lymphoid enhancer-binding factor 1 |
| 22 | MIMAT0010251 | hsa-miR-449c-5p | CCNG1 | Cyclin G1 |
| 23 | MIMAT0019780 | hsa-miR-4690-3p | PKM | Pyruvate kinase M1/M2 |
| 24 | MIMAT0019880 | hsa-miR-4746-5p | LMO2 | LIM domain only protein 2 |
| 25 | MIMAT0019904 | hsa-miR-4758-3p | MMP16 | Matrix metalloproteinase-16 |
| 26 | MIMAT0002883 | hsa-miR-514a-3p | RBX1 | Ring-box 1 (ROC1) |
| 27 | MIMAT0002868 | hsa-miR-522-3p | PDK1 | Pyruvate dehydrogenase kinase isoform 1 |
| 28 | MIMAT0003260 | hsa-miR-592 | PIK3CA | Phosphatidylinositol-4,5-bisphosphate 3-kinase catalytic subunit alpha |
| 29 | MIMAT0025450 | hsa-miR-6499-5p | ARHGAP23 | Rho GTPase-activating protein 23 |
| 30 | MIMAT0027467 | hsa-miR-6783-3p | ITGA5 | Integrin alpha-5 |
| 31 | MIMAT0028118 | hsa-miR-7110-3p | PDGFA | Platelet-derived growth factor subunit A |
| 32 | MIMAT0028125 | hsa-miR-7114-5p | RUNX1T | RUNX1 partner transcriptional co-repressor 1 |
| 33 | MIMAT0004945 | hsa-miR-744-5p | ATXN2 | Ataxin-2 |
| 34 | MIMAT0004957 | hsa-miR-760 | HIST2H3C | Histone H3.2 |
| 35 | MIMAT0031177 | hsa-miR-7974 | CREB3L3 | Cyclic AMP-responsive element-binding protein 3-like protein 3 |
| 36 | MIMAT0004950 | hsa-miR-877-3p | MMP16 | Matrix metalloproteinase-16 |
| 37 | MIMAT0004902 | hsa-miR-891a-5p | EN1 | Engrailed homeobox 1 |
| 38 | MIMAT0004907 | hsa-miR-892a | ZEB1 | Zinc finger E-box-binding homeobox 1 (TCF8) |
| 39 | MIMAT0004980 | hsa-miR-937-3p | PTGFRN | Prostaglandin F2 receptor inhibitor (TIR8 or SIGIRR) |
| 40 | MIMAT0004983 | hsa-miR-940 | IGF1 | Insulin-like growth factor 1 |

**Table 5B:**
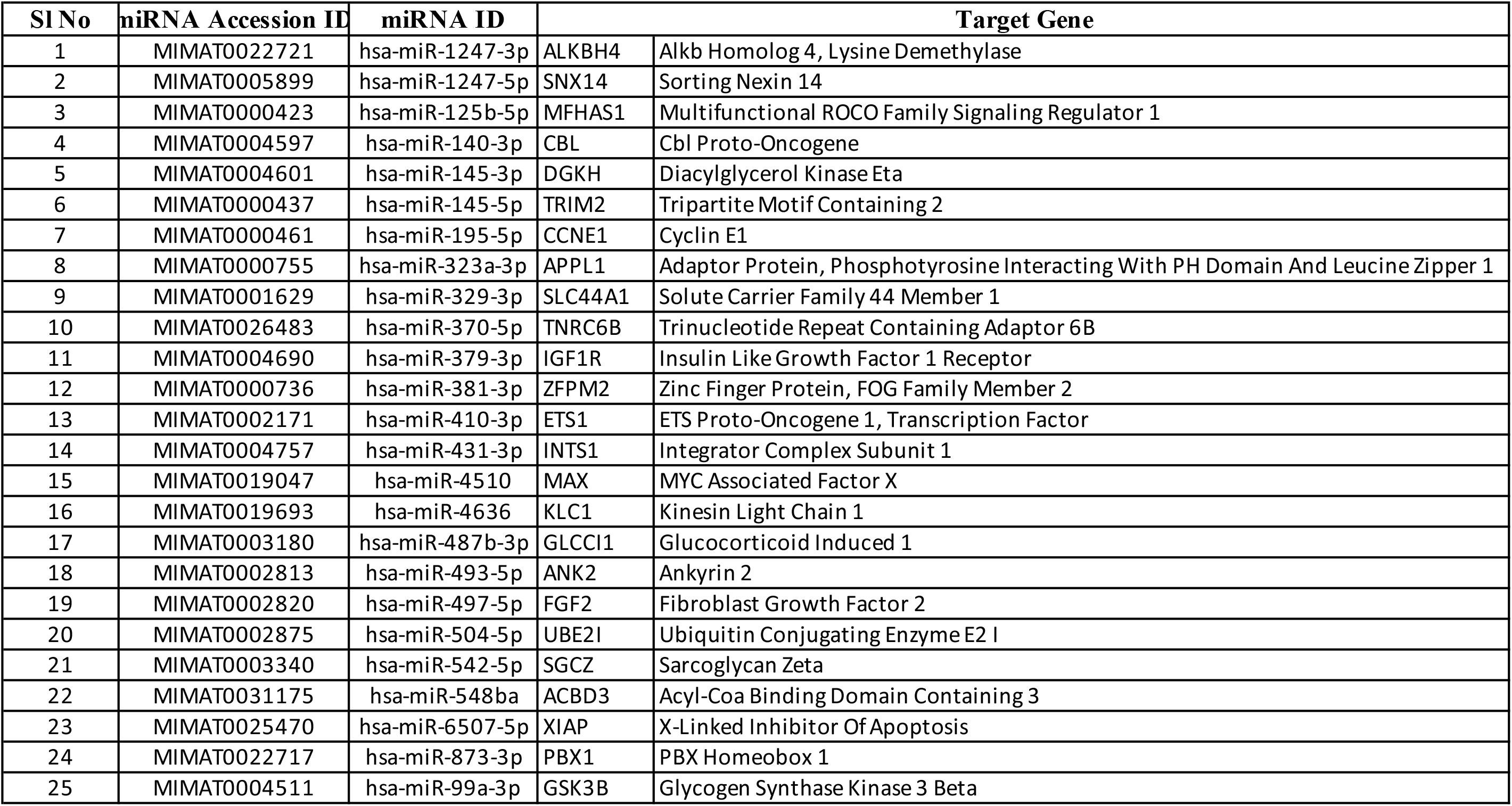
List of predicted mRNA targets for the underexpressed miRNAs which have significant impact on Overall survival.

| Sl No | miRNA Accession ID | miRNA ID | Target Gene |  |
| --- | --- | --- | --- | --- |
| 1 | MIMAT0022721 | hsa-miR-1247-3p | ALKBH4 | Alkb Homolog 4, Lysine Demethylase |
| 2 | MIMAT0005899 | hsa-miR-1247-5p | SNX14 | Sorting Nexin 14 |
| 3 | MIMAT0000423 | hsa-miR-125b-5p | MFHAS1 | Multifunctional ROCO Family Signaling Regulator 1 |
| 4 | MIMAT0004597 | hsa-miR-140-3p | CBL | Cbl Proto-Oncogene |
| 5 | MIMAT0004601 | hsa-miR-145-3p | DGKH | Diacylglycerol Kinase Eta |
| 6 | MIMAT0000437 | hsa-miR-145-5p | TRIM2 | Tripartite Motif Containing 2 |
| 7 | MIMAT0000461 | hsa-miR-195-5p | CCNE1 | Cyclin E1 |
| 8 | MIMAT0000755 | hsa-miR-323a-3p | APPL1 | Adaptor Protein, Phosphotyrosine Interacting With PH Domain And Leucine Zipper 1 |
| 9 | MIMAT0001629 | hsa-miR-329-3p | SLC44A1 | Solute Carrier Family 44 Member 1 |
| 10 | MIMAT0026483 | hsa-miR-370-5p | TNRC6B | Trinucleotide Repeat Containing Adaptor 6B |
| 11 | MIMAT0004690 | hsa-miR-379-3p | IGF1R | Insulin Like Growth Factor 1 Receptor |
| 12 | MIMAT0000736 | hsa-miR-381-3p | ZFPM2 | Zinc Finger Protein, FOG Family Member 2 |
| 13 | MIMAT0002171 | hsa-miR-410-3p | ETS1 | ETS Proto-Oncogene 1, Transcription Factor |
| 14 | MIMAT0004757 | hsa-miR-431-3p | INTS1 | Integrator Complex Subunit 1 |
| 15 | MIMAT0019047 | hsa-miR-4510 | MAX | MYC Associated Factor X |
| 16 | MIMAT0019693 | hsa-miR-4636 | KLC1 | Kinesin Light Chain 1 |
| 17 | MIMAT0003180 | hsa-miR-487b-3p | GLCCI1 | Glucocorticoid Induced 1 |
| 18 | MIMAT0002813 | hsa-miR-493-5p | ANK2 | Ankyrin 2 |
| 19 | MIMAT0002820 | hsa-miR-497-5p | FGF2 | Fibroblast Growth Factor 2 |
| 20 | MIMAT0002875 | hsa-miR-504-5p | UBE2I | Ubiquitin Conjugating Enzyme E2 I |
| 21 | MIMAT0003340 | hsa-miR-542-5p | SGCZ | Sarcoglycan Zeta |
| 22 | MIMAT0031175 | hsa-miR-548ba | ACBD3 | Acyl-Coa Binding Domain Containing 3 |
| 23 | MIMAT0025470 | hsa-miR-6507-5p | XIAP | X-Linked Inhibitor Of Apoptosis |
| 24 | MIMAT0022717 | hsa-miR-873-3p | PBX1 | PBX Homeobox 1 |
| 25 | MIMAT0004511 | hsa-miR-99a-3p | GSK3B | Glycogen Synthase Kinase 3 Beta |

**Table 6:**
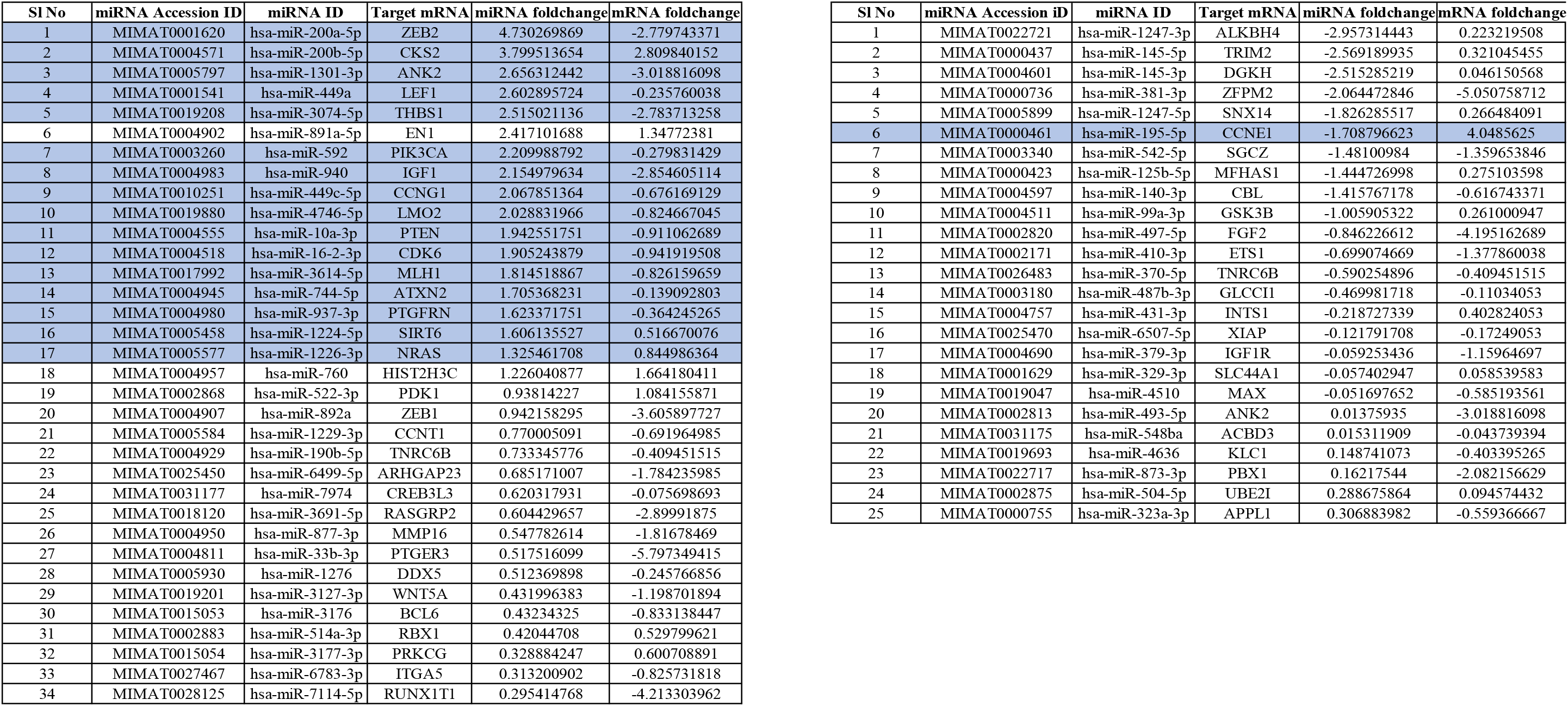
List of miRNA-mRNA pairs which could be potential biomarkers for prognosis.

## Discussion

Endometrial cancer is the 6^th^ most common cancer in women worldwide and the 15^th^ most common cancer worldwide (43). If diagnosed early (stage 1 or 2), patients with endometrial cancer have a good prognosis, while patients diagnosed at later stages don’t respond well to chemotherapy and have poor outcomes. The categorization of patients as early stage or late stage is mainly based on histopathological evaluation mostly post-surgery. However, this is not a very reliable method in endometrial cancer since it has been seen that cancers of the same stage and histology can have very different molecular profiles (reviewed in (3)). Moreover, classification based on histopathology would be subject to variability between observers, which in many cases can lead to lapses in patient care, either under or over treatment. Therefore, classification/staging requires better molecular markers so that therapeutic approaches can be better strategized.

Currently, markers such as mutations in POLE, MMR and p53 are being used for endometrial cancer classification (19). However, these tests require a combination of histopathology and molecular techniques and hence may not be very feasible in routine clinical settings. Studies on multiple cancers including endometrial cancer have identified miRNAs as good diagnostic and prognostic markers (44). miRNAs have good potential to be diagnostic and prognostic markers since it is seen that the miRNAs expression is cell-type specific, de-regulated in many disease conditions, and also because they can be analyzed with samples procured with non-invasive procedures (14). Based on studies in other cancers such as breast cancer, miRNAs are good markers for early and sensitive diagnosis of cancers (45).

In the context of endometrial cancer, miRNA expression data is available from other populations but not from Indian population. Our data shows a miRNA profile from Indian patients. We have identified 195 differentially expressed miRNAs in endometrial cancers as compared to control samples. Further, we have compared our miRNA profile with that available from TCGA and found that 169 miRNAs were common to Indian population and the other populations on TCGA (Figure 1, Table 3). Since ours is a limited time study, we did not have survival data for the Indian population. Hence, the differential expression data from this population has been correlated to survival data from TCGA to get a miRNA signature relevant in prognosis. From this analysis, we have identified 65 miRNAs in endometrial cancer whose expression levels may have a relevance in overall survival of patients. We have also identified potential mRNA targets for these miRNAs. This information could further be used to understand the mechanism of action of the miRNAs and how they may affect the behavior of the tumor cells. Also, this study has identified 59 miRNA-mRNA pairs which are relevant in endometrial cancer. Out of these, 16 miRNA-mRNA pairs could be used as potential biomarkers with prognostic relevance (Table 6). There have been some studies which have identified differentially expressed miRNAs in endometrial cancer and their potential role in prognosis. Using archival FFPE samples, 84 cancer specific miRNAs have been identified as differentially expressed in endometrial cancers. Some of these miRNAs also have an implication in prognosis (21). When this data is compared with our cohort, only 2 miRNAs (hsa-miR-195-5p and hsa-miR-125b-5p) were in common. This could be due to the fact that the above study was PCR based and analyzed a limited number of miRNAs and our study being NGS based could identify even novel miRNAs.

Another study that has performed a meta-analysis of miRNA data from TCGA has identified some potential miRNAs as well as miRNA-mRNA pairs that could have prognostic significance in endometrial cancer (23). A study conducted at Ohio State University Medical Center, Columbus, OH, from January 1997–July 2003 has done a microRNA profiling of surgically staged endometrial cancers. They have identified differentially expressed miRNA between normal and tumor and also some miRNAs like miR199 which have an implication in patient outcome (22). Although these two studies are similar to our study, they have analyzed patients from one particular region. Our data represents population of a different ethnicity, which is not very well represented in other studies including TCGA.

The major limitation of our study is that the profiling has been done on one sub-type of endometrial cancer as identified by histopathology. Also, the samples collected here are prospective samples collected between 2018 and 2022 and hence we do not have survival data for these patients. Despite these limitations, the approach used here as summarized in figure 4 is comprehensive. This is the first reported miRNA profile from Indian population. The comparison and correlation with data from TCGA makes it more globally relevant since it considers a larger and more diverse population. Further, correlation to overall survival makes the list of biomarkers significant in prognosis. This approach can also be extended across sub-types of endometrial cancer. This kind of molecular-subtyping would be very valuable in deciding the best course of treatment for the patients. The 16 miRNA-mRNA pairs as identified by this study, if validated across a larger sample size, has the potential to be developed into a very useful diagnostic tool.

**Figure 4:**
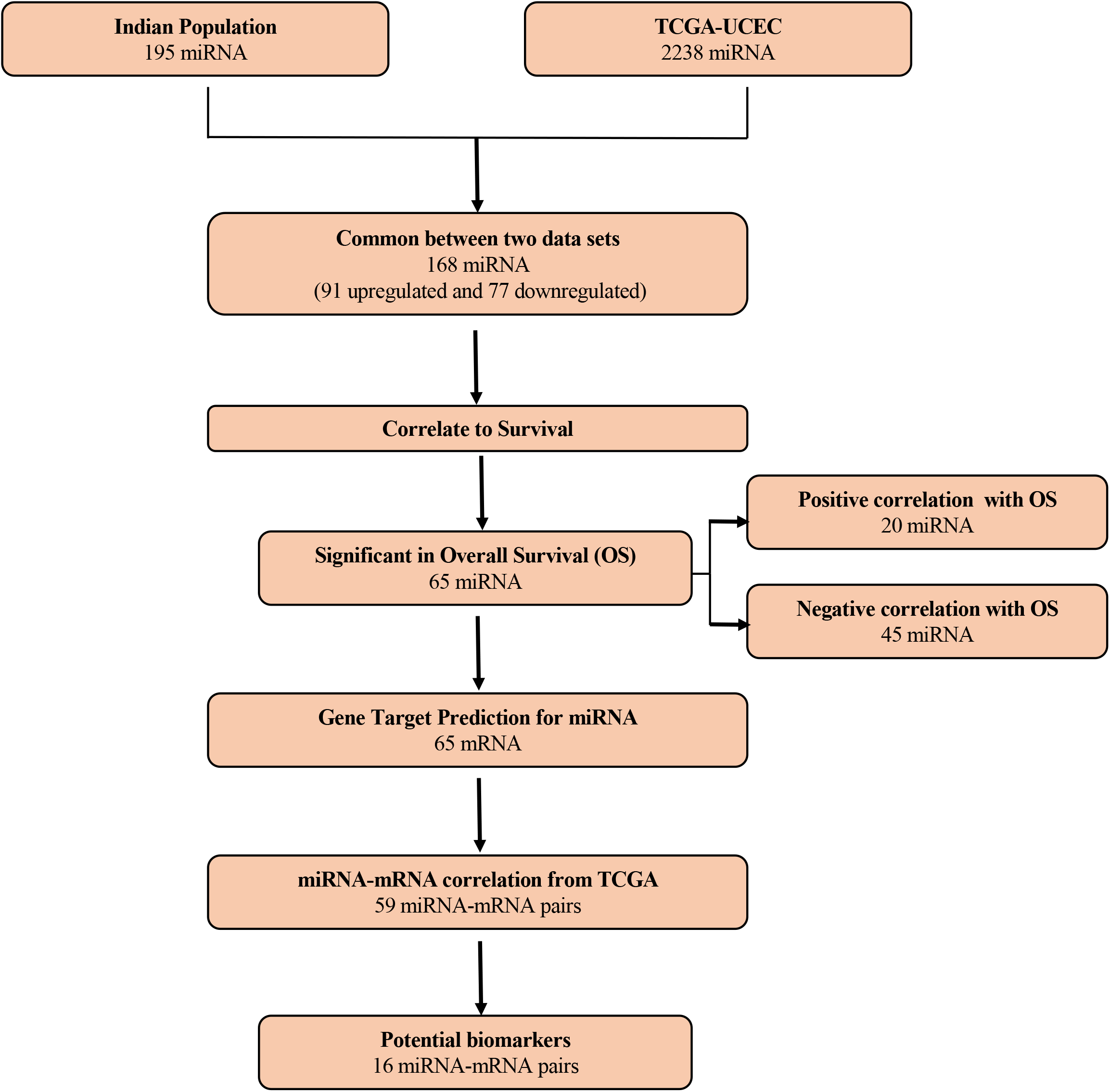
Schematic work flow for identification of potential biomarkers for endometrial cancer

## Declaration

The work described here has been approved by the IEC of all the participating institutions, namely KMIO, VV and CHG, Bengaluru, India.

This work has not been published anywhere yet.

## Author Contributions

The project was conceived and strategized by PR and SMN; Sample procurement and collection clinical details was done by PVR, SC and SMN. Sample collection was coordinated by SA and NHN, Sample processing and inventory was done by SN, Data analysis has been performed by SH, KW and AA. Manuscript was written by PR and assisted by KW and AA. All authors have contributed to the manuscript and have approved of it.

SH, KW and AA have contributed equally for this manuscript.

## Figure and table legends

**Table 1:** List of differentially expressed miRNAs between control endometrium and endometrial cancer. The list shows the miRNA ID, log2 fold change and the p value.

1A: List of miRNAs overexpressed in endometrial cancer as compared to control tissue (114)

1B: List of miRNAs under-expressed in endometrial cancer as compared to control tissue (81)

**Table 2:** List of differentially expressed novel miRNAs in endometrial cancer. The table shows the log2 fold change, the consensus sequence, chromosomal location and provisional ID of these miRNAs.

**Table 3:** List of common miRNAs between TCGA data set and Indian data

3A: List of common miRNAs overexpressed in endometrial cancer (91)

3B: List of common miRNAs under-expressed in endometrial cancer (77)

**Table 4:** List of differentially expressed miRNAs which have significant impact on overall survival

4A: miRNAs whose expression correlates positively with survival (20)

4B: miRNAs whose expression correlates negatively with survival (45)

**Table 5:** List of predicted mRNA targets for the miRNAs which have significant impact on overall survival

5A: mRNA targets for overexpressed miRNAs (40)

5B: mRNA targets for under-expressed miRNAs (25)

**Table 6:** List of miRNA-mRNA pairs which could be potential biomarkers for prognosis

Note: The table shows the log2 fold changes of the miRNA and the target mRNA according to TCGA data. The highlighted cells are the pairs which show inverse correlation between miRNA and the corresponding mRNA target.

Note: Only those miRNAs which show similar pattern of expression in Indian data and TCGA data have been considered.

## Acknowledgements

We thank all members of PR lab for their support during this study and preparation of manuscript. Special thanks to Dr. Dixcy Jaba Sheeba and Namratha Nadig for helping out during the initial stages of the project.

